# Lung epithelial signaling mediates early vaccine-induced CD4^+^ T cell activation and *Mtb* control

**DOI:** 10.1101/2021.06.03.446981

**Authors:** Shibali Das, Nancy D. Marin, Ekaterina Esaulova, Mushtaq Ahmed, Amanda Swain, Bruce A. Rosa, Makedonka Mitreva, Javier Rangel-Moreno, Mihai G Netea, Luis B. Barreiro, Maziar Divangahi, Maxim N. Artyomov, Deepak Kaushal, Shabaana A. Khader

## Abstract

Tuberculosis (TB) is one of the leading causes of death due to a single infectious agent. The development of a TB vaccine that induces durable and effective immunity to *Mycobacterium tuberculosis* (*Mtb*) infection is urgently needed. Early and superior *Mtb* control can be induced in *M. bovis* Bacillus Calmette–Guérin (BCG) vaccinated hosts when the innate immune response is targeted to generate effective vaccine-induced immunity. In the present study, we show that innate activation of DCs is critical for mucosal localization of clonally activated vaccine-induced CD4^+^ T cells in the lung, and superior early *Mtb* control. In addition, our study reveals that Th1/Th17 cytokine axis play an important role in superior vaccine induced immunity. Our studies also show that activation of nuclear factor kappa-light-chain-enhancer of activated B cells (NFκβ) pathway in lung epithelial cells is critical for the mucosal localization of activated vaccine-induced CD4^+^ T cells for rapid *Mtb* control. Thus, our study provides novel insights into the immune mechanisms that can overcome TB vaccine bottlenecks and provide early rapid *Mtb* control.

**Importance:** Tuberculosis is a leading cause of death due to single infectious agent accounting 1.4 million deaths each year. The only licensed vaccine BCG is not effective due to variable efficacy. In our study, we determined the early immune events necessary for achieving complete protection in BCG vaccinated host. Our study reveals that innate activation of DCs can mediate superior and early *Mtb* control in BCG vaccinated host through lung epithelial cell signaling and localization of clonal activated, *Mtb* antigen specific, cytokine producing CD4^+^ T cells within the lung parenchyma and airways. Thus, our study provides novel insights into the immune mechanisms that can overcome TB vaccine bottlenecks and provide early rapid *Mtb* control.

## Introduction

*Mycobacterium tuberculosis* (*Mtb*) is a leading cause of death worldwide by a single infectious agent and it infects approximately one fourth of the world’s population (1). Currently, *M. bovis* Bacillus Calmette-Guerin (BCG) is the only licensed vaccine against tuberculosis (TB). However, the variable efficacy of BCG, along with the emergence of drug resistant *Mtb* strains and comorbidity associated with Human Immunodeficiency Virus infection, has further confounded the eradication of TB as a public health problem. Recently, the M72/AS01E vaccine has been reported to provide about 50% efficacy in *Mtb*-infected adults against pulmonary TB disease (2). Additionally, use of BCG vaccination either mucosally or intravenously (IV) in rhesus macaques substantially limited *Mtb* infection (3–5). However, despite these break-through findings in the TB vaccine field, we do not fully understand the mechanistic basis behind the early immune events within the lungs that mediates protection in vaccinated hosts.

T cells are required to control *Mtb* in vivo as mice lacking CD4^+^ T cells are unable to control *Mtb* infection (6). *Mtb* infection is known to delay initiation of the adaptive T cell immune response resulting in early uncontrolled *Mtb* growth (7–9). Moreover, during *Mtb* infection, the colocalization of immune cells, including CD4^+^ T cells and macrophages within the lung parenchyma rather than in the lung vasculature is critical for early *Mtb* control (10, 11). However, the exact mechanisms that mediate the recruitment of CD4^+^ T cells in vaccinated hosts to mediate *Mtb* control is unclear. In the current study, using a mouse model of early superior *Mtb* control in BCG vaccinated hosts, we have delineated the early protective mechanisms that mediate vaccine-induced *Mtb* control. We demonstrate that the localization of clonally expanded *Mtb*-specific cytokine-producing CD4^+^ T cell population which preferentially localize in the lung parenchyma and airways, are critical for early *Mtb* control. Additionally, early signaling events in lung epithelial cells are critical to facilitate the interaction between activated *Mtb*-specific CD4^+^ T cells and macrophages, within the parenchyma and airways, for rapid *Mtb* control in this model. Thus, our study provides novel immunological insights into the early mechanism of vaccine-induced protective immunity against TB, allowing for potential targeting these pathways to improve TB vaccine efficacy for future use.

## Results

### Innate activation of DCs amplifies clonal vaccine-induced CD4^+^ T cell responses in *Mtb*- infected BCG vaccinated host

Delayed activation and accumulation of *Mtb-*specific vaccine-induced T cells in the lung is a critical bottleneck for vaccine protection against *Mtb* infection (8, 9). Innate activation by transfer of exogenously zymosan activated *Mtb* antigen 85B (Ag85B)-pulsed DCs (Z-DC) into BCG vaccinated *Mtb*-infected hosts resulted in early CD4^+^ T cell recruitment, enhanced IFN-γ and IL-17 production and complete early control of *Mtb* replication in mice (8). Using this published model of early and superior vaccine-induced *Mtb* control, we probed the exact immune mechanism(s) of protection in vaccinated hosts. The peak of the CD4^+^T cell response in unvaccinated *Mtb*-infected mice is 20 dpi, while the peak of the vaccine response is 15 dpi following *Mtb* infection (12). We also showed that in Vac+Z-DC group, the vaccine responses are accelerated at 8 and 15 dpi (8). Therefore, we picked the peak time point as the appropriate measurement of effective T cell responses between the conditions in order to assess their full expression of antimycobacterial function. As a first step, lung cells were isolated at the peak of the immune response respectively from *Mtb*-infected unvaccinated C57BL/6 mice at 20 days post infection [dpi] (Unvac), *Mtb*-infected BCG vaccinated C57BL/6 mice at 15 dpi (Vac), and *Mtb*-infected BCG vaccinated C57BL/6 mice that received Z-DC transfer (given intratracheally at -1 and +4 dpi) at 8 and 15 dpi (Vac+Z-DC) (**Fig.1A**) and subjected to single cell RNA sequencing (scRNA-Seq) to define the immune cell populations driving enhanced vaccine-induced immunity.

**Figure 1.**
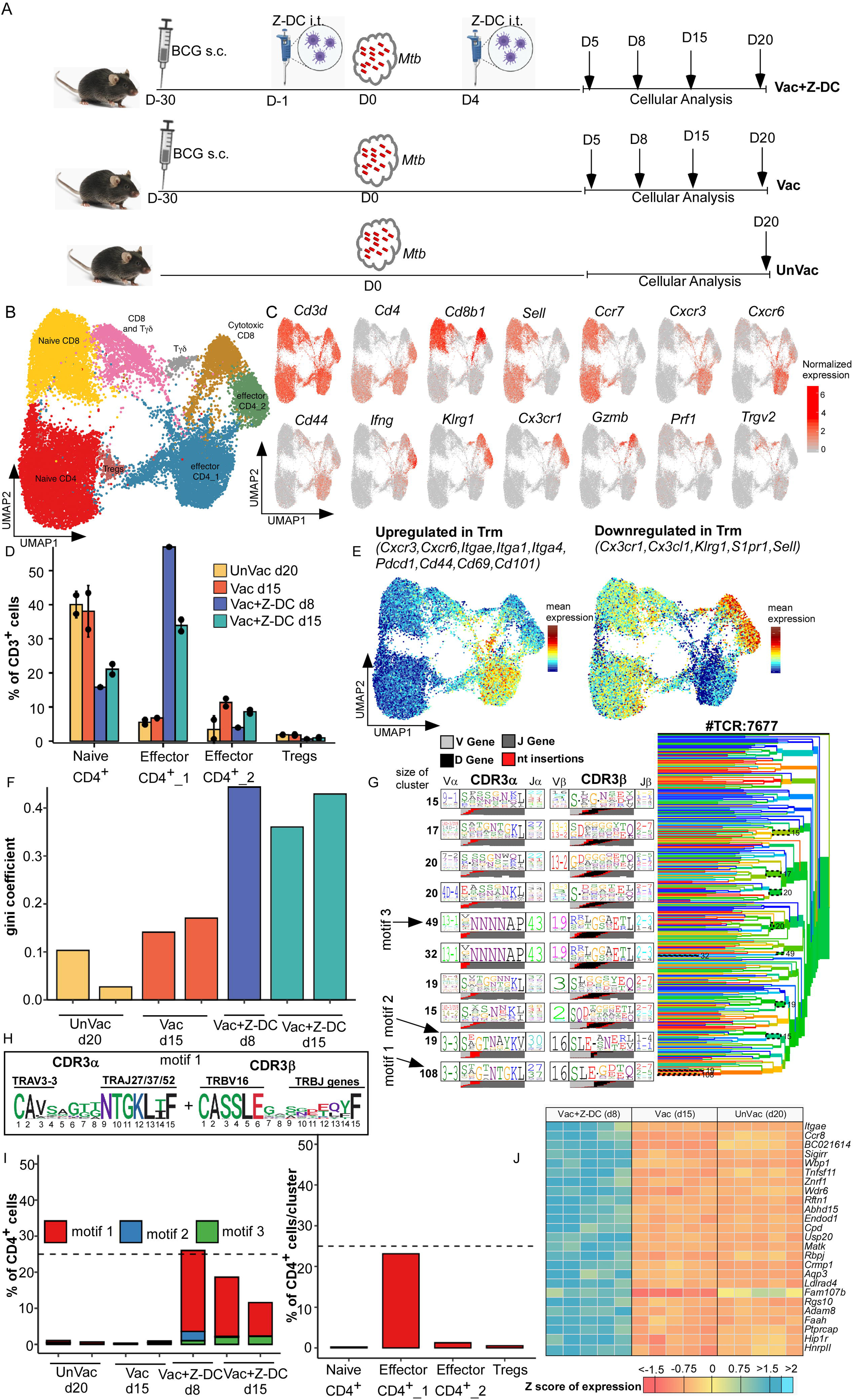
Activation of DCs amplifies rapid CD4^+^ T cell responses in BCG vaccinated *Mtb* infected mice. C57BL/6 (B6) mice were either left unvaccinated (UnVac) or vaccinated with BCG (Vac), rested for 4 weeks and infected with *Mtb* HN878. Some BCG vaccinated mice received Z-DC (Vac+Z-DC) at the time of *Mtb* infection (A). Lungs were harvested at different dpi and single cell suspensions were subjected to scRNA-Seq. UMAP with lung isolated CD3^+^ cells, combined plot shown from unvaccinated (20 dpi) (n=2), vaccinated (15 dpi) (n=2), Vac+Z-DC (8 and 15 dpi) (n=1 and n=2 respectively) conditions are presented here (**B**). UMAPs with marker genes used to assign identity to clusters of T cells are shown (**C**). CD4^+^ cluster abundances as percent of total CD3^+^ cells across four conditions are shown. Error bars are mean ± SD for two replicates from each condition. Vac+Z-DC 8 dpi had only one replicate (**D**). Mean expression of genes, upregulated (left panel) or downregulated (right panel) in lung T resident memory (Trm) cells on UMAP for all conditions are shown (**E**). Gini coefficient for CD4^+^ repertoires across all samples are shown (**F**). TCRdist tree result for 7677 unique CD4^+^ TCRs are shown (**G**). Amino acid composition of CDR3α and CDR3β for motif 1 depicted as sequence logo (**H**). Proportion of CD4^+^ cells, matching motif 1 across all samples (left panel) and across CD4^+^ clusters (right panel) are shown (**I**). Gene expression profile of total CD4^+^ T cells isolated from *Mtb*-infected unvaccinated (20 dpi), *Mtb*-infected BCG vaccinated (15 dpi), and *Mtb*- infected BCG vaccinated C57BL/6 mice that receive Z-DC transfer (8 dpi), was determined by RNA sequencing **(J).** Z-scores calculated from the FPKM values across all of the samples, individually for each gene. n = 5 biological replicates for gene expression profile analysis.

In CD3^+^ cells, unsupervised clustering yielded four clusters of CD4^+^ cells, namely naive CD4^+^ (*Ccr7^+^Sell^+^*), two clusters of effector CD4^+^ cells (*Ccr7^-^Sell^-^CD44^+^* or *Ifng^+^*) and T regulatory cells (Foxp3^+^); three clusters of CD8^+^ cells - naive CD8^+^ (*Ccr7^+^Sell^+^*), cytotoxic CD8^+^ (*Gzmb^+^Prf1^+^*), mix of CD8^+^ and Tγδ cells; and more enriched Tγδ cluster (*Trgv2^+^*) (**Fig. 1B, C and Fig. S1A, Table S1**). Notably, cluster of naive CD4^+^ T cells dominated in both *Mtb*-infected unvaccinated and BCG vaccinated mice and decreased in *Mtb*-infected BCG vaccinated mice that received Z-DC transfer (40% versus 20% of T cells). In contrast, effector CD4_1 cluster dominated the T cell population in *Mtb*-infected BCG vaccinated mice that received Z-DC transfer (harvested at either 8 or 15 dpi) while being almost absent in *Mtb*-infected BCG vaccinated or unvaccinated mice (35-55% versus <10% of total T cells, **Fig. 1D and S1B**). Effector CD4_1 exhibited the signature of lung T resident memory (Trm) cells including *Cxcr3, Cxcr6, Itgae, Cd44, Cd69 and Cd101*(13) (**Fig. 1E and Fig. S1C**). We have used the publically available surface receptor markers of Trm cells in lungs of humans as broad lung T cell tissue resident signature (13, 14). Gene set enrichment analysis (GSEA) shows an enrichment of genes, upregulated in lung Trm (GSE94964), in our comparison between CD4_1 and CD4_2/Naive CD4 clusters. Incidentally, we also identified TCR for each cell, and first compared clonotype expansion with Gini coefficient (coefficient of 0 means that each T cell clonotype has only 1 T cell, while coefficient of 1 means that all T cells have identical clonotype). CD4^+^ cells from *Mtb-*infected BCG vaccinated mice that received Z-DC transfer had Gini coefficient twice higher than *Mtb*-infected unvaccinated or BCG vaccinated mice, suggesting that *Mtb-*infected BCG vaccinated mice that received Z-DC transfer have more expanded clonotypes, when compared to CD4^+^ T cells in *Mtb*-infected unvaccinated or BCG vaccinated mice (**Fig. 1F**).

We also identified potential motif-based groups of CD4^+^ T cells recognizing the same epitopes derived from antigens among the TCRs using TCRdist tool. We focused on the 3 most well-defined motifs with conserved amino acids, coming from TCR clusters of size 108, 19 and 49 TCRs (**Fig. 1G,H, Fig S1D and Table S2**). As TCRdist operates on unique TCR sequences without taking into account clonotype expansion, we enumerated the number of CD4^+^ T cells expressing any of these TCR motifs, and we found that motif 1 was exclusively associated with *Mtb*-infected BCG vaccinated mice that received Z-DC transfer, where it was present in 10-25% of all CD4^+^ T cells (**Fig. 1I**, left panel **and Fig. S1E**), while other motifs were much less expanded. Motif 1 is characterized by TRAV3-3 - TRAJ27/37/52 - TRBV16 pairing, and it features sub-motifs in both CDR3α and CDR3β. Conserved amino acids come from V and J gene parts rather than from insertion of random nucleotides. Furthermore, we found that majority of cells with motif 1 belonged to effector CD4_1 cluster (**Fig. 1I,** right panel), accounting for >20% of cells inside the cluster. We next analyzed transcriptional differences among the CD4^+^ T cells between different groups. Therefore, we isolated the CD4^+^ T cells (at the time of peak CD4^+^ T cell responses) from *Mtb*-infected unvaccinated C57BL/6 mice at 20 dpi, *Mtb*-infected vaccinated C57BL/6 mice at 15 dpi and *Mtb*-infected vaccinated C57BL/6 mice that received Z-DC transfer at 8 dpi (**Fig. 1J and Fig. S2A**). Among the top 25 differently expressed genes in the *Mtb*-infected BCG vaccinated mice that received Z-DC transfer, there was a marked upregulation of genes associated with T cell migration including *Ccr8* (*15*)*, Itgae (Cd103)* (16)*, Aqp3* (17) and *Rbpj* associated with cell-cell communications (18). Surprisingly, gene expression profile of CD4^+^ T cells from *Mtb*-infected unvaccinated or BCG vaccinated mice were comparable (**Fig. 1J**, **Fig. S2A and B**). The important enriched pathways associated with T cells receptor signaling and T cell function were observed in the *Mtb*-infected BCG vaccinated mice that received Z-DC transfer compared with the CD4^+^ T cells isolated from BCG vaccinated mice (**Fig. S2C**). Therefore, our data suggest that CD4^+^ T cells that confer early *Mtb* control in *Mtb*-infected BCG vaccinated mice that received Z-DC transfer express functionally distinct T cell transcriptional profiles associated with migration.

### Early activation and mucosal localization of CD4^+^ T cells mediates improved *Mtb* control in BCG vaccinated host

We next determined the functional ability of the vaccine-induced CD4^+^ T cells to promote macrophage killing of *Mtb*. We isolated highly pure lung CD4^+^ T cells from either *Mtb*-infected BCG vaccinated C57BL/6 mice (at 15 dpi) or *Mtb*-infected BCG vaccinated C57BL/6 mice that received Z-DC transfer (at 8 dpi) and co-cultured isolated CD4^+^ T cells with *Mtb*-infected macrophages in vitro to assess *Mtb* killing. Interestingly, both CD4^+^ T cells from *Mtb*-infected BCG vaccinated mice and *Mtb*-infected BCG vaccinated mice that received Z-DC transfer effectively mediated comparable *Mtb* killing (**Fig. 2A**). Additionally, while IFN-γ production was similar in supernatants from co-cultures of both groups that received CD4^+^ T cells (**Fig. 2B**), higher levels of IL-17 were detected in co-cultures that received CD4^+^ T cells from *Mtb*-infected BCG vaccinated mice that received Z-DC transfer, when compared to co-cultures that received *Mtb*-infected BCG vaccinated CD4^+^ T cells (**Fig. 2C**). As we did not find any functional differences in *Mtb* killing within macrophages between the two groups, we hypothesized that the clonally expanded CD4^+^ T effector population was mediating improved protection possibly due to other mechanisms rather than just direct activation of macrophages. Thus, we next isolated highly pure lung CD4^+^T cells from *Mtb*-infected vaccinated mice that received Z-DC transfer (at 8 dpi) and adoptively transferred the CD4^+^ T cells into the BCG vaccinated C57BL/6 mice following *Mtb* infection, while control mice were BCG vaccinated and *Mtb*-infected that did not receive T cells. The rationale was to test if adoptive transfer improved *Mtb* control when compared with just BCG vaccination. Adoptive transfer of purified CD4^+^ T cells into *Mtb*-infected BCG vaccinated mice resulted in improved *Mtb* control, when compared to PBS treated *Mtb*-infected BCG vaccinated mice (**Fig. 2D**), and this coincided with improved B cell follicle formation associated with immune control of *Mtb* (19), without impacting overall lung inflammation (**Fig. 2E and F**). This improved *Mtb* control coincided with dampened production of proinflammatory cytokines in the lungs of Mtb-infected BCG vaccinated mice that received CD4^+^ T cells including IL-12, TNF-α, IL-10, IL-1β and IL-6, and the chemokines KC, MIP-1β, RANTES and MIP-2 (**Fig. 2G and H**) corroborating with the unaltered overall lung inflammatory landscape. Together, these results suggest that vaccine-induced CD4^+^ T cell drive protection in *Mtb*-infected BCG vaccinated hosts likely by their ability to migrate and localize into specific lung compartments. Therefore, we next studied the kinetics associated with the CD4^+^ T cell activation as well as localization of CD4^+^ T cells in the lung of *Mtb*-infected BCG vaccinated mice and *Mtb*-infected BCG vaccinated mice that received Z-DC transfer. Remarkable CD4^+^ T cell activation (CD44^hi^) was observed as early as 3 dpi (**gating strategy in Fig. S2D)** in *Mtb*-infected BCG vaccinated mice that received Z-DC transfer, and this correlated with significant and rapid accumulation of *Mtb* Ag85B tetramer-specific (TET^+^) CD4^+^ T cells (within the lungs of *Mtb*-infected BCG vaccinated mice that received Z-DC transfer compared to *Mtb*-infected BCG vaccinated mice **(Fig. 3A, and Fig. S3A**). These robust and early responses were maintained, with 8 dpi being the peak of the response. We observed nearly 400-fold higher CD4^+^CD44^hi^TET^+^ T cells in *Mtb*-infected BCG vaccinated mice that received Z-DC transfer, when compared with *Mtb*-infected BCG vaccinated mice. In contrast, the expansion and accumulation of CD4^+^CD44^hi^TET^+^ T cells in *Mtb*-infected BCG vaccinated mice was delayed until 20 dpi (**Fig. 3A).**

**Figure 2.**
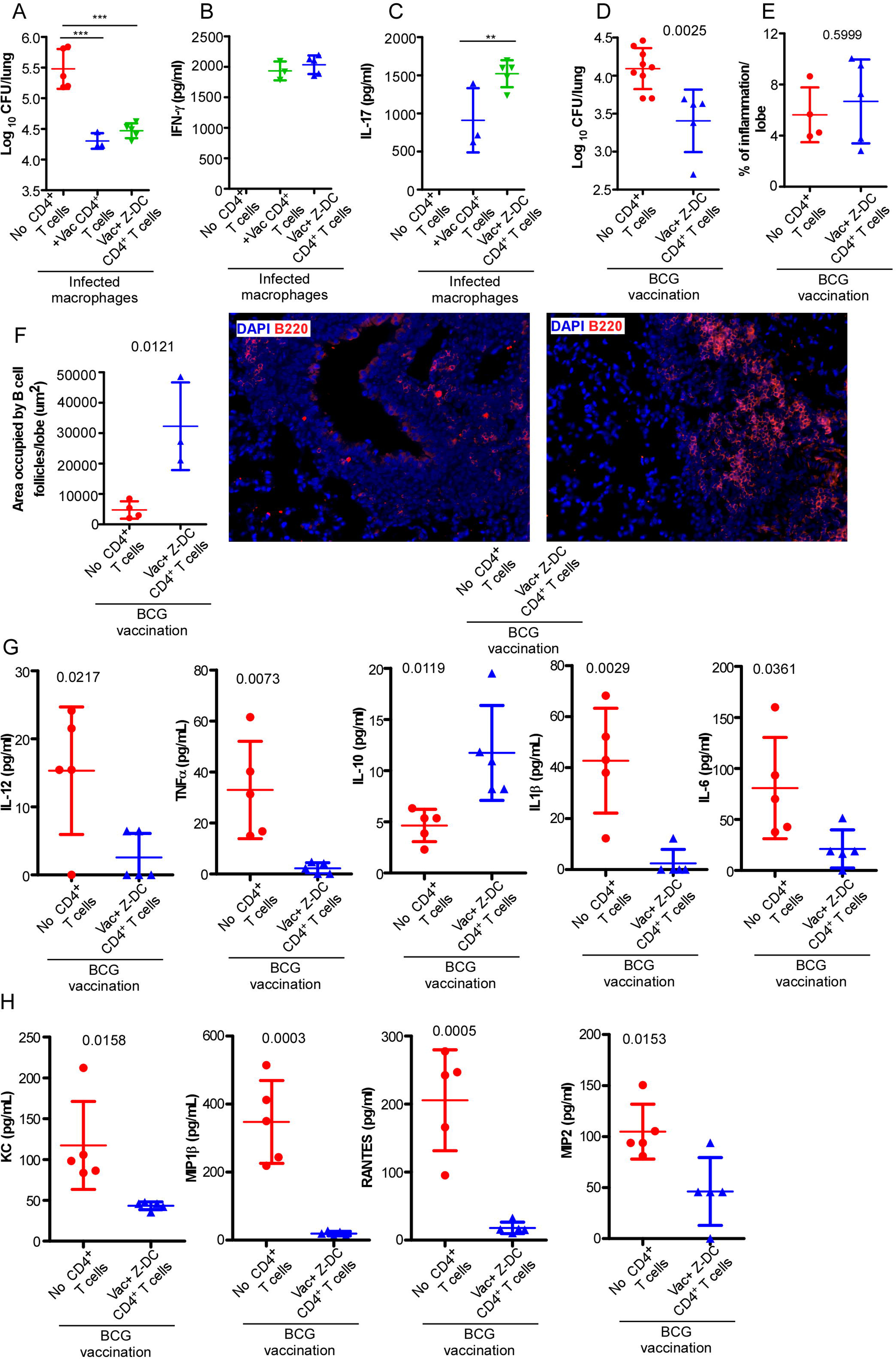
Adoptive transfer of vaccine-induced CD4^+^ T cells mediates improved *Mtb* control in BCG vaccinated host. CD4^+^ T cells were isolated from *Mtb*-infected BCG vaccinated C57BL/6 mice and *Mtb*-infected BCG vaccinated C57BL/6 mice that received Z-DC transfer and co-cultured with *Mtb*-infected BMDMs (1:1 ratio) for six days. Intracellular *Mtb* CFU was determined by plating cell lysates **(A)**. IFN-γ and IL-17 levels **(B, C)** were measured in cell supernatants by ELISA. n = 3-5 biological replicates. C57BL/6 mice were vaccinated with BCG, rested for 4 weeks and infected with *Mtb* HN878. CD4^+^ T cells were isolated from *Mtb*-infected BCG vaccinated mice that received Z-DC transfer (at 8 dpi) and adoptively transferred to *Mtb*-infected BCG vaccinated mice. Lungs were harvested at 30 dpi and lung bacterial burden was determined by plating **(D).** Lung inflammation was calculated in the H&E stained FFPE lung sections (**E**). B cell lymphoid follicle formation was determined on the FFPE lung sections by B220 (red) immunofluorescence staining (**F**). n = 3-9 mice per group. Levels of cytokines (**G**) and chemokines (**H)** in lung homogenates were quantified by multiplex. n = 5 biological replicates. Data represented as mean ± SD. ND = not detected. ** p≤0.01, *** p≤0.001 either by one way ANOVA (**A-C)** or by Student’s t test (actual p values are shown) **(D-H)**.

**Figure 3.**
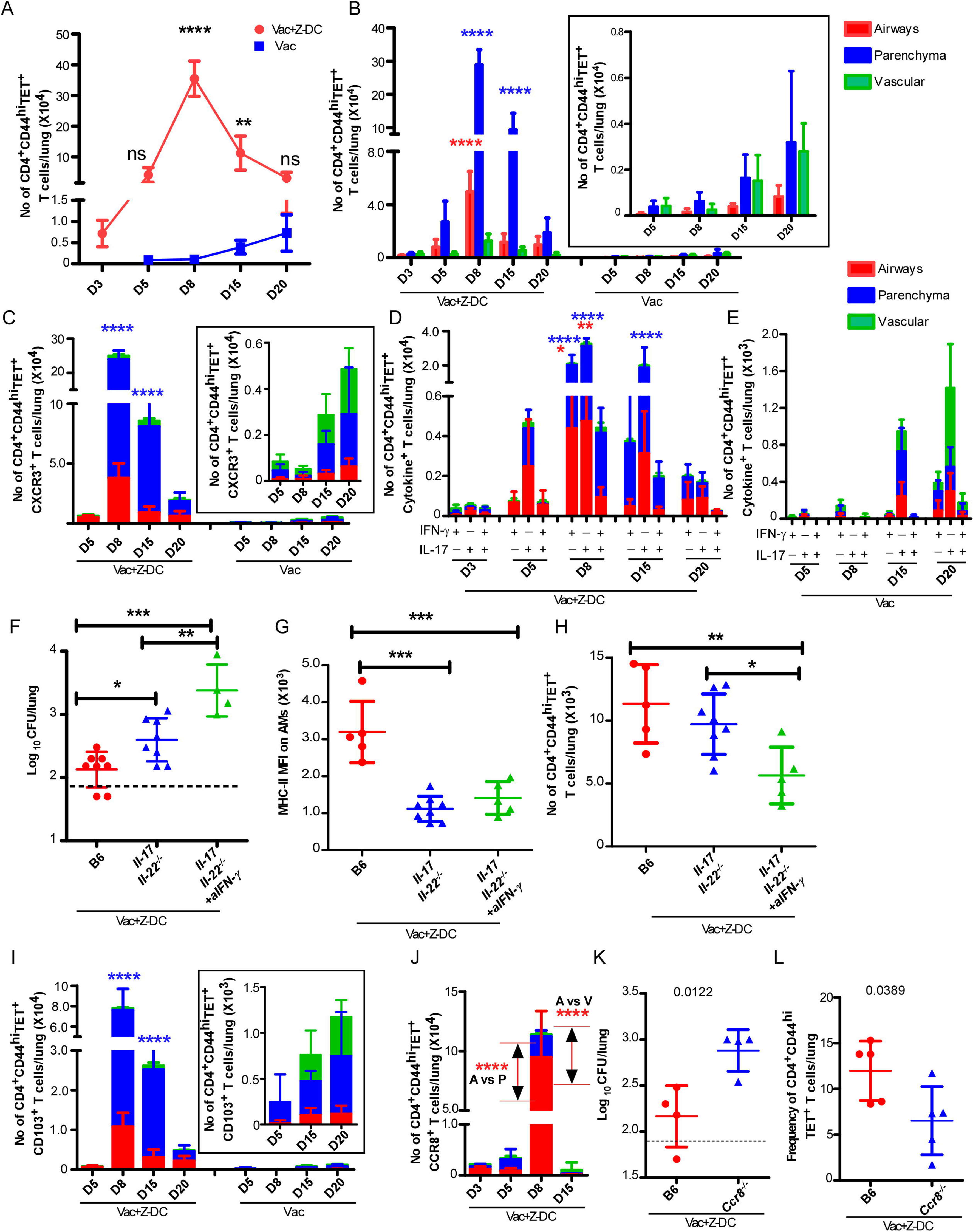
Rapid and early CD4^+^ T cell activation and localization within lung parenchyma and airway is driven by CCR8 engagement. C57BL/6 mice were vaccinated, *Mtb*-infected and received Z-DC transfer as described in method. At the time of harvest, the mice received anti-CD45.2-v500 (IT) and anti-CD45.2-BV605 (IV) antibodies as described in method. Lungs were harvested and subjected to flow cytometry to measure the number of CD4^+^CD44^hi^TET^+^ (**A**) T cells. The distribution of CD4^+^CD44^hi^TET^+^ (**B**), CD4^+^CD44^hi^TET^+^CXCR3^+^ (**C**), CD4^+^CD44^hi^TET^+^cytokine^+^ (**D, E**), CD4^+^CD44^hi^TET^+^CD103^+^ (**I**), CD4^+^CD44^hi^TET^+^CCR8^+^ (**J**) T cells in groups of BCG vaccinated mice were measured in lung airways (red bar), parenchyma (blue bar) and vasculature (green bar) regions by flow cytometry. n=4-5 mice per group. B6, and *IL-17/IL-22^-/-^* mice were vaccinated with BCG, rested for 4 weeks and infected with *Mtb* HN878 and received Z-DC. One group of BCG vaccinated *Mtb*-infected *IL-17/IL-22^-/-^* mice received IFN-γ neutralizing ab. Mice were harvested at 20 dpi and lung bacterial burden was determined by plating **(F)**. MHC-II MFI (mean fluorescent intensity-MFI) on AMs (**G**) and number of CD4^+^CD44^hi^TET^+^ T cells **(H)** were assessed by flow cytometry on total lung single cell suspensions. n=4-5 mice per group. B6, and *Ccr8^-/-^* mice were vaccinated with BCG, rested for 4 weeks and infected with *Mtb* HN878. Some BCG vaccinated mice received Z-DC. Mice were harvested at 20 dpi and lung bacterial burden was determined by plating **(K)**. Frequency of CD4^+^CD44^hi^TET^+^ T cells **(L)** were assessed by flow cytometry on total lung single cell suspensions. n=4-8 mice per group. Data represented as mean + SD.* p≤0.05, ** p≤ 0.01, *** p≤ 0.001, ****p≤ 0.0001 either by two-way ANOVA (**A-E, I and J**), one-way ANOVA (**F-H**) or Student’s t test (actual p values are shown) (**K and L**).

Immune cell recruitment in airways and parenchyma correlated with increased protection compared to localization in lung vasculature during *Mtb* infection (11, 20). To identify the localization of activated T cells to specific lung compartments, we tracked vasculature-localization (CD45.2-BV605^+^), airway localization (CD45.2-V500^+^) and parenchyma localization (BV605 negative V500 negative) of immune cells in the lung (10, 11, 21) by administering anti-CD45.2-BV605 intravascularly or anti-CD45.2-V500 intratracheally just prior to sacrifice. We found that CD4^+^CD44^hi^ (**Fig. S3B**), CD4^+^CD44^hi^TET^+^ T cells (**Fig. 3B**) localized to the lung parenchyma of *Mtb-*infected BCG vaccinated mice that received Z-DC transfer at very early time points (3-15 dpi), with progressive accumulation of CD4^+^CD44^hi^ and CD4^+^CD44^hi^TET^+^ T cells into the airways over time. Coincident with the delayed accumulation of CD4^+^CD44^hi^TET^+^ T cells in *Mtb*-infected BCG vaccinated mice, the majority of the CD4^+^CD44^hi^TET^+^ T cells localized mainly in vasculature and parenchyma with some cells localizing within the airways (**Fig. 3B, inset**). These data together suggest that activation of the innate immune pathways to target vaccine-induced T cell responses can initiate rapid expansion of CD4^+^CD44^hi^TET^+^ T cells with specific localization in the lung parenchyma and airways, contributing to early and rapid *Mtb* control.

CXCR3 is a well-described chemokine receptor expressed by circulating T cells. As infection progressed, the accumulation of CD4^+^CD44^hi^TET^+^CXCR3^+^ T cells increased into the lung parenchyma and airways in *Mtb-*infected BCG vaccinated mice that received Z-DC transfer (**Fig. 3C)**. However, reduced accumulation of CD4^+^CD44^hi^TET^+^CXCR3^+^ T cells in the lung parenchyma and airways in *Mtb-*infected BCG vaccinated mice was observed (**Fig. 3C, inset)**. The CD4^+^CD44^hi^TET^+^CXCR3^+^ cells which expanded at d15 post infection exhibited a ratio of parenchyma: vasculature associated T cells of 16.69 (± 10.54) in Vac+Z-DC mice as compared with BCG vaccinated mice (3.794 ± 6.572, p 0.0489 by Student’s T test between Vac+Z-DC and BCG vaccinated *Mtb*-infected mice). IFN-γ and IL-17 are important effector cytokines that contribute to protective immune responses against *Mtb* infection (3, 22, 23). While the majority of the CD4^+^CD44^hi^TET^+^ T cells were IL-17^+^ cytokine-producing, a population of IFN-γ^+^ cytokine-producing and IFN-γ^+^ /IL-17^+^ double cytokine-producing CD4^+^CD44^hi^TET^+^ T cells were also found in the lungs of *Mtb*-infected vaccinated mice that received Z-DC transfer and they accumulated as early as 5 dpi with peak responses at 8 dpi (**Fig. 3D**). Importantly, the cytokine-producing CD4^+^CD44^hi^TET^+^ T cells predominantly localized to the lung parenchyma and airways. Moreover, in *Mtb*-infected BCG vaccinated mice that received Z-DC transfer, the frequency of IL-17^+^ cytokine-producing CD4^+^CD44^hi^TET^+^ T cells showed an early and sustained increase, when compared with IFN-γ^+^ and IFN-γ^+^IL-17^+^ cytokine-producing CD4^+^CD44^hi^TET^+^ T cells. Indeed, consistent with delayed accumulation of CD4^+^CD44^hi^TET^+^ T cells in *Mtb*-infected BCG vaccinated lungs, the number of IFNγ^+^ and IL-17^+^ single or dual cytokine-producing CD4^+^CD44^hi^TET^+^ T cells were delayed and 10 fold lower in *Mtb*-infected BCG vaccinated mice, and they preferentially localized in the vasculature (**Fig. 3E**). To fully characterize the role of cytokine signaling in Z-DC mediated protection in the BCG vaccinated *Mtb* infected mice, we transferred Z-DC in BCG vaccinated IL-17/IL-22 double knockout (*Il-17/Il-22^-/-^*) mice with or without IFN-γ neutralization to evaluate the specific contribution of Th1 and/or Th17 responses. Absence of IL-17/IL-22 signaling together led to significantly higher bacterial burden (**Fig. 3F**) as compared with the wild type BCG vaccinated C57BL/6 mice, which also received Z-DC transfer. These results suggest an important role for IL-17/IL-22 signaling, specifically under conditions of the superior protection enabled by Z-DC transfer in BCG vaccinated mice. Additionally, we observed reduced expression of MHC-II expression on alveolar macrophages (AMs) (**Fig. 3G, gating strategy in Fig. S3C**) within the lung of BCG vaccinated *Il-17/Il-22^-/-^* mice receiving Z-DC transfer. Moreover, blocking IFN-γ in BCG vaccinated *Il-17/Il-22^-/-^* mice which also received Z-DC transfer, abrogates *Mtb* control with significant reduction in expression of MHC Class II expression on AMs and reduced accumulation of CD4^+^CD44^hi^TET^+^ T cells, compared with the wild type BCG vaccinated C57BL/6 mice receiving Z-DC transfer (**Fig. 3G,H**). Therefore, our study points towards a synergistic role played by the Th1-Th17 axis in mediating better protection in the BCG vaccinated *Mtb* infected mice receiving Z-DC. These results suggest that the protection mediated in the *Mtb*-infected BCG vaccinated mice that received Z-DC transfer is associated with an early activation, expansion and localization of cytokine-producing CD4^+^ T cells within the parenchyma and airways where *Mtb*-infected macrophages are harbored.

Both scRNA-Seq and bulk gene signature analysis of CD4^+^ T cells demonstrated elevated expression of genes associated with migration such as *Itgae* or the genes expressed by tissue resident T cells such as the chemokine receptor, *Ccr8* (**Fig. 1B and I**). Expression of CD103 and CCR8 on CD4^+^CD44^hi^TET^+^ T cells peaked on 8 and 15 dpi respectively which corresponded with the enhanced recruitment of CD4^+^CD44^hi^TET^+^ T cells into lungs and the early control of *Mtb* replication in BCG vaccinated mice that received Z-DC transfer (**Fig. 3I, J and insets**). In addition, *Mtb*-infected *Cd103* deficient BCG vaccinated mice that received Z-DC transfer exhibited similar protective capacity than *Mtb*-infected wild type mice that also received Z-DC transfer (**Fig. S3D**), suggesting that CD103 expression did not induce additional protective mechanisms in BCG vaccinated mice that received Z-DC transfer. In contrast, *Ccr8* deficient *Mtb*-infected BCG vaccinated mice receiving Z-DC transfer did not control *Mtb* replication to the similar level as observed in wild type *Mtb*-infected BCG vaccinated mice that received Z-DC transfer (**Fig. 3K**). This decreased vaccine-induced control in *Ccr8* deficient *Mtb*-infected BCG vaccinated mice receiving Z-DC transfer coincided with reduced accumulation of CD4^+^CD44^hi^TET^+^ T cells within the lung of *Ccr8* deficient mice (**Fig. 3L**). These results suggest CD103-independent but CCR8-dependent mechanisms underlying Z-DC mediated vaccine-induced *Mtb* control. However, it could also be possible that CCR8 along with CD103 synergistically regulate the CD4^+^ T cell localization and activation to impact *Mtb* replication in vivo and needs further experimentation.

### Lung epithelial signaling is critical for early immune cell activation and mucosal localization in *Mtb*-infected vaccinated mice

Our data demonstrate that mucosal localization of CD4^+^ T cells within the parenchyma and airways is effective at inducing complete early *Mtb* control in vaccinated hosts. Our recent studies showed that AMs upon activation migrate from the airways into the parenchyma to form granulomas and mediate effective *Mtb* control (10). Thus, we next addressed if localization of CD4^+^ T cells in the parenchyma and airways resulted in more effective and early activation of AMs. We observed that AMs readily accumulated in the lung airways and parenchyma of *Mtb-* infected BCG vaccinated mice that received Z-DC transfer (**Fig. 4A, gating strategy in Fig.S3C)** with increasing accumulation and rapid early upregulation of MHC Class II expression as an indicator of activation **(Fig. 4B)**. In contrast, AM localization within the airways in the *Mtb*- infected BCG vaccinated mice lungs was delayed with fewer AMs that accumulated within the parenchyma and delayed timing of AM activation. In BCG vaccinated mice, while there is no significant accumulation of AMs in the parenchyma, there is a small but significant increase in AM population within the airways at 15 dpi, when compared to 8 dpi. Additionally, we observed a marked early recruitment and activation of recruited macrophages (RMs), mainly in airways and lung parenchyma of the *Mtb*-infected BCG vaccinated mice that received Z-DC transfer. In contrast, in the *Mtb*-infected BCG vaccinated mice, RMs localized mainly in vasculature and at lesser extent in parenchyma (**Fig. 4C and D**). Thus, collectively our data suggest that early infiltration and localization of myeloid cells within specific lung parenchyma are crucial factors for inducing superior vaccine-induced immunity.

**Figure 4.**
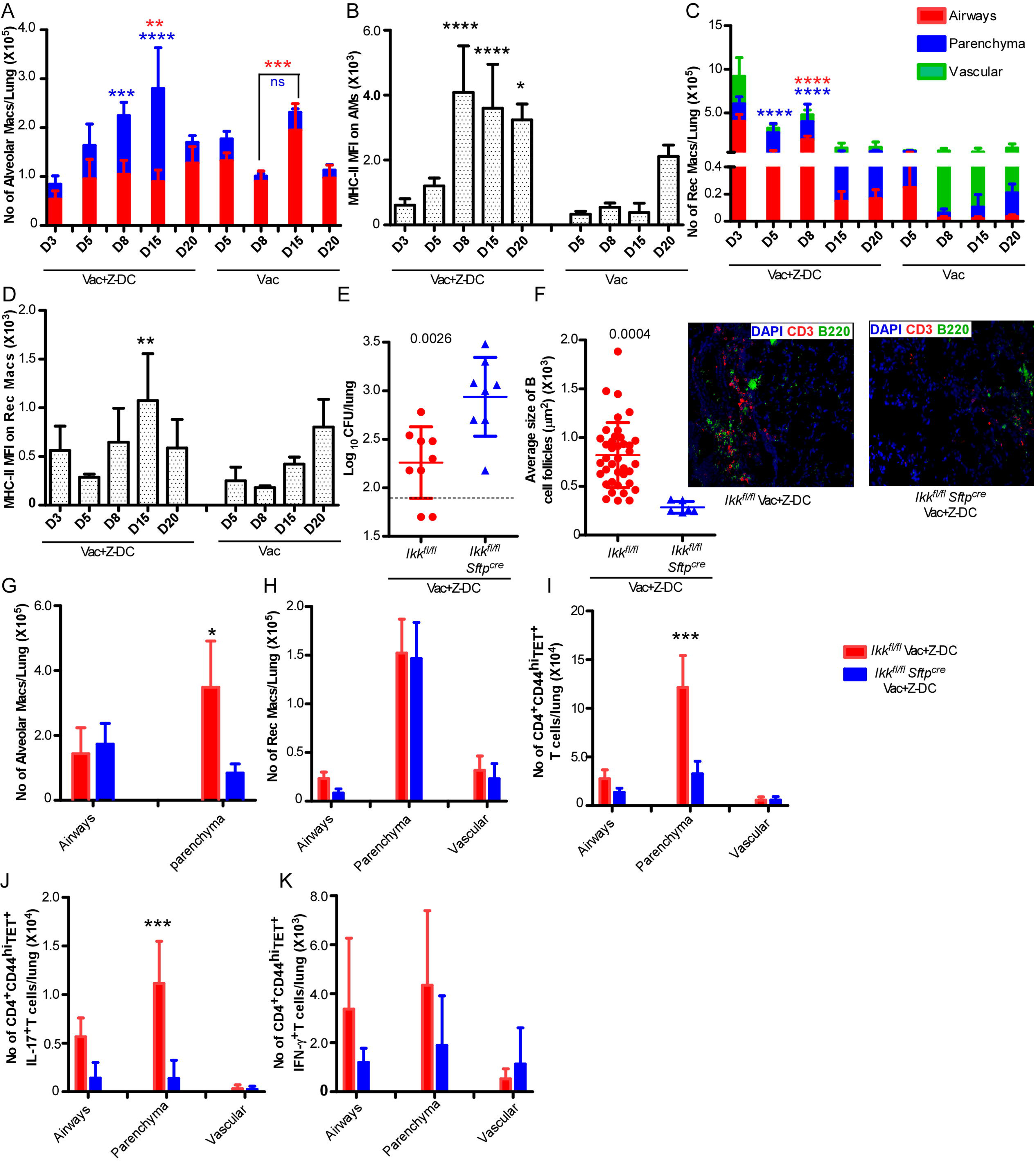
NFκβ signaling in lung epithelial cells mediates early CD4^+^ T cell activation and mucosal localization. C57BL/6 mice were vaccinated, *Mtb*-infected and received Z-DCs as described in method. To track the myeloid cells, mice were given anti-CD45.2-v500 and anti-CD45.2-BV605 antibodies through IT and IV route respectively prior to harvest. Lungs were harvested at different dpi and total numbers of AMs **(A)**,MHC-II MFI (mean fluorescent intensity-MFI) on AMs **(B)**, RMs **(C)**, MHC-II MFI on RMs **(D)** in airways (red bar), parenchyma (blue bar) and vasculature (green bar) location were assessed by flow cytometry. n=4-5 mice per group. In a separate experiment, *Ikk*^fl/fl^ and *Ikk^fl/fl^Sftpcre* mice were vaccinated with BCG, infected and received Z-DCs as described under method. To track the immune cells, mice were given anti-CD45.2-v500 and anti-CD45.2-BV605 antibodies through IT and IV route respectively prior to harvest. Lungs were harvested at 20 dpi and lung bacterial burden was determined by plating **(E)**. B cell lymphoid follicles were determined by CD3 (red) and B220 (green) staining on FFPE lung sections by immunofluorescence staining (**F**). The total numbers of AMs **(G)**, RMs **(H),** CD4^+^CD44^hi^TET^+^ (**I**), CD4^+^CD44^hi^TET^+^cytokine^+^ (**J, K)** T cells in the airways, parenchyma and vasculature location were determined by flow cytometry (red=*Ikk^fl/fl^*, n=4-9; blue= *Ikk^fl/fl^Sftpcre*, n=4-8). Data represented as mean ± SD.* p≤ 0.05, ** p≤ 0.01, *** p≤ 0.001, ****p≤ 0.0001 either by two way ANOVA (**A-D, G-K**) or by Student’s t test (actual p values are shown) (**E and F**).

Epithelial cells respond to IL-17/IL-22 (24–26) and secrete various antimicrobial peptides and several chemokines (27), that orchestrate recruitment of immune cells such as neutrophils, recruited monocytes and natural Th17 cells (28, 29) . Thus, we hypothesized that activation of epithelial signaling will participate in production of mediators involved in strategic immune cell localization and early control of *Mtb* infection. To further delineate mechanistic insight of the signaling pathways associated with the superior vaccine-induced immunity in *Mtb*-infected BCG vaccinated mice that received Z-DC transfer, we used *Ikk^fl/fl^Sftpcre* mice which lack NFκΒ signaling specifically in lung type II epithelial cells and therefore do not upregulate the expression of necessary inflammatory cytokines or chemokines (10, 30). Absence of functional lung epithelial signaling in *Mtb-infected* BCG vaccinated *Ikk^fl/fl^Sftpcre* mice that received Z-DC transfer did not provide the superior and early vaccine-induced *Mtb* control as compared with the *Mtb*-infected BCG vaccinated *Ikk^fl/fl^* littermate control mice that received Z-DC transfer (**Fig. 4E**). These data suggest that functional epithelial signaling plays an important role in inducing the superior protection specifically under conditions of Z-DC transfer into BCG vaccinated mice following *Mtb*-infection. Moreover, we observed reduced B cell follicle formation within the lungs of *Mtb*-infected BCG vaccinated *Ikk^fl/fl^Sftpcre* mice that received Z-DC transfer compared to *Mtb*- infected BCG vaccinated littermate control mice that received Z-DC transfer (**Fig. 4F)**. To further investigate the influence of functional epithelial signaling on immune mechanisms, we studied the immune cell localization in *Mtb*-infected BCG vaccinated *Ikk^fl/fl^Sftpcre* and littermate control mice that received Z-DC transfer. As expected, we observed reduced AM accumulation within lung parenchyma in *Mtb-*infected BCG vaccinated *Ikk^fl/fl^Sftpcre* mice that received Z-DC transfer as compared with *Mtb*-infected BCG vaccinated littermate controls mice that received Z-DC transfer (**Fig. 4G)**. Although we did not see any defects in the accumulation of RMs in *Mtb*- infected BCG vaccinated *Ikk^fl/fl^Sftpcre* mice that received Z-DC transfer (**Fig. 4H)**, we observed defective activation of RMs in *Mtb*-infected BCG vaccinated *Ikk^fl/fl^Sftpcre* mice that received Z-DC transfer as compared with *Mtb*-infected BCG vaccinated littermate control mice that received Z-DC transfer (**Fig. S3E)**. Moreover, we found a significant reduction in the number of CD4^+^CD44^hi^ and CD4^+^CD44^hi^TET^+^ T cells present in lung parenchyma of *Mtb*-infected BCG vaccinated *Ikk^fl/fl^Sftpcre* mice that received Z-DC transfer as compared with *Mtb*-infected BCG vaccinated littermate control mice that received Z-DC transfer (**Fig. S3F,G and Fig. 4I**). Similarly, the number of CD4^+^CD44^hi^TET^+^IL-17^+^ cytokine-producing T cells present in lung parenchyma of *Mtb*-infected BCG vaccinated *Ikk^fl/fl^Sftpcre* mice that received Z-DC transfer was significantly reduced compared with *Mtb*-infected BCG vaccinated littermate control mice that received Z-DC transfer (**Fig. 4J**). The number of CD4^+^CD44^hi^TET^+^IFN-γ^+^ cytokine-producing CD4^+^ T cells present in lung airways and parenchyma of *Mtb*-infected BCG vaccinated *Ikk^fl/fl^Sftpcre* mice that received Z-DC transfer was lower, as compared with *Mtb*-infected BCG vaccinated littermate control mice that received Z-DC transfer although it was not statistically significant (**Fig. 4K**). Thus, our data suggest that activation of lung epithelial signaling regulates the environmental signals that mediate localization and amplification of Th17/Th1 responses within the site of infection to mediate early superior control of *Mtb* infection in the vaccinated host.

## Discussion

The development of a TB vaccine that induces durable and effective immunity to *Mtb* infection is urgently needed. Previously, we have demonstrated that complete and early *Mtb* control can be induced in BCG vaccinated hosts when the innate immune response is targeted to generate effective vaccine-induced immunity. In the present study, we show that the mechanisms by which activation of innate immunity results in superior *Mtb* control is through rapid and robust amplification of cytokine-producing vaccine-induced T cell responses and localization within the airways and parenchyma of BCG vaccinated hosts. Our studies also show that activation of NFκβ pathway in lung epithelial cells is an important early event that drives the localization of vaccine-induced CD4^+^ vaccine-induced T cells within parenchyma and activation of myeloid cells, thus promoting the formation of protective iBALT structures within the lung and control of *Mtb* replication. Therefore, our study provides novel insights into the immune mechanisms that can overcome TB vaccine bottlenecks and provide early rapid *Mtb* control.

*Mtb* is a successful pathogen due to its ability to evade host immune responses. Studies have shown that following *Mtb* infection, delay in the activation of antigen specific CD4^+^ T cell responses occurs likely due to *Mtb*’s ability to directly inhibit MHC-II trans activator expression, MHC–II expression and antigen presentation (31). BCG vaccination can generate systemic vaccine-induced T cell responses, but upon *Mtb* challenge, the accumulation of T cells to the lung although accelerated when compared with naïve hosts (day 15 when compared to day 20 in naïve), is still not sufficiently early or durable enough to provide complete protection against *Mtb* infection (8, 32). Therefore, as shown in our previous work(8) and by others (33), targeting the innate pathway through DC activation is one way to rapidly activate T cell responses to mediate early and rapid control. Our new studies shown here, demonstrate that the mechanisms by which innate DC activation overcomes the roadblock is by rapid amplification of early CD4^+^ T cell responses by 5 days post *Mtb* challenge, and localization within airway and parenchyma compartments in the lung. That these CD4^+^ T cells are recruited and localized within 5 days, is by far the earliest recruitment of vaccine-induced T cells responses to most vaccine strategies against TB. This early amplification is similar to what is seen in even after 30 days following BCG IV vaccination in NHPs where the heightened and lung-localized Trm cells are considered to be a mechanism through which complete control of *Mtb* infection is mediated (3). In the current study, we show that adoptive transfer of peptide pulsed Z-DC into BCG vaccinated mice provides superior *Mtb* control. We have previously shown that following BCG vaccination (34), the accumulation of antigen-specific lung-resident cytokine-producing T cells in the lung is not as robust as amplification of antigen-specific T cells in the spleen and lymph nodes of vaccinated mice. Based on these data, we expect that adoptive transfer of Z-DC results in activation of antigen-specific T cells in the lymph nodes and possibly recruitment to the lung. However, we don’t rule out the possibility of local proliferation of T cells following Z-DC transfer. Thus, our results along with recent studies including IV (3) and mucosal BCG use (5), suggest that overcoming the *Mtb* suppression-mediated of early immune responses is thus possible and generating lung-resident activated T cell pool should be a good strategy for improving vaccine-induced immunity against TB.

During *Mtb* infection, Ag85B is predominantly secreted during the early phase of infection and expression reduced by 3 weeks post infection, while ESAT-6 is expressed and secreted by *Mtb* during chronic phases of infection (35). Since we are interested in the early events post infection, we designed our study to activate innate immune responses to amplify BCG-vaccine induced Ag85B specific CD4^+^ T cell responses against *Mtb* infection. Indeed, our results show that early amplification of Ag85B specific T cell responses resulted in complete control of *Mtb* infection. As several vaccine formulations including whole vaccines such as BCG and subunit vaccine such as H56/IC31 (36) and H56/CAF01 (37) include Ag85B as an antigen, our results suggest that targeting innate DC activation to rapidly amplify Ag85B-specific CD4^+^ T cell responses may further improve upon protection elicited by these vaccines in animal models and should be robustly tested. Furthermore, our studies for the first time demonstrate that in fact the transcriptional profiles that are induced in BCG vaccine-induced CD4^+^ T cell responses are comparable to CD4^+^ T cells induced in unvaccinated *Mtb*-infected mice, suggesting that the quality of responses induced by BCG vaccination is not very different from *Mtb* infection. This is in sharp contrast to the early amplification of vaccine-induced CD4^+^ T cells found in BCG vaccinated hosts that undergo innate DC activation where there was a marked upregulation of genes associated with T cell migration and cell-cell communications, especially expression of genes such as Cd103 and Ccr8 which likely allow localization of effector T cells into lung compartment for optimal *Mtb* control. Thus, our results also provide novel insights that both the timing of arrival of CD4^+^ T cells and the quality (cytokine production and upregulation of receptors and proteins enabling lung localization) of CD4^+^ T cells are important for optimal *Mtb* control, compared to CD4^+^ T cells induced by BCG vaccination that are not fully protective.

For the first time as far as we are aware, our study was able to identify potential motif-based groups of CD4^+^ T cells recognizing the epitopes of same antigens among the TCRs. Our results showed that a motif was exclusively associated with T cells in *Mtb*-infected BCG vaccinated mice that received Z-DC transfer, where it was present in 10-25% of all CD4^+^ T cells. These results suggest that innate DC activation of T cells along with amplification of CD4^+^ T cell also allows for clonal expansion of activated Trm cells. Previous studies have demonstrated that in mice vaccinated with Ag85B had a skewed CDR3β length distribution with preferential use of TRBV16 and two CDR3βs namely CASSLEGDEQYF and CASSLEGDTQYF (32). Our studies have not only validated the presence of the CDR3β motifs but also identified the motif on CDR3α that are amplified in CD4^+^ T cells from BCG vaccinated mice that also received Ag85B primed Z-DCs. Our studies identified 3 CDR3α and 3 CDR3β motifs highly represented in effector CD4^+^ T cells from BCG vaccinated mice that also received Ag85B primed Z-DCs. From the available literature(38) the predominant Motif 1 appears to be specific for *Mtb* Ag85B. The presence of “LEG” motif in the TCR sequence specifically in TCRβ, identifies the TCR specific for Ag85B. Carpenter et al, 2017 showed the TCRβ repertoire of vaccine-elicited (Ag85B _240-254_) and *Mtb*-recalled Ag85b-specific CD4^+^ T cells, as well as after primary infection.

Therefore, unlike the unvaccinated or BCG vaccine induced TCR repertoire on CD4^+^ T cells, Z-DC induced TCR repertoire demonstrated in this study represents novel motifs which have potential to control *Mtb* infection. Thus, while bulk RNA-seq studies do not allow us due to the pooled nature of cells to understand the heterogeneity of T cell responses, scRNA-Seq allows us to understand the heterogeneity of T cell responses. For example, our results show that while activated population of clonally expanded CD4^+^ T cells expressing Trm markers increase in BCG vaccinated host receiving Z-DC transfer, this population is not different between BCG vaccinated and unvaccinated lungs. Instead, an IFN-γ^+^ CD4^+^ T cell population is increased in BCG vaccinated lungs when compared with unvaccinated lungs. Therefore, our results highlight the utility of using single cell TCR sequencing to probe the expansion and clonality of vaccine-responsive CD4^+^ T cells and provide an in-depth understanding of T cell responses generated following vaccination.

During *Mtb* infection, effective control of intracellular *Mtb* requires direct recognition of infected macrophages in the lung by CD4^+^ effector T cells (39). Therefore, localization of Trm CD4^+^ T cells in the lung is an important event required for *Mtb* control. CXCR3 expression on CD4^+^ T cells (CXCR3^+^) is considered a marker of lung recruited CD4^+^ T cells and is important for localization of CXCR3^+^ Th1 cells to the lung parenchyma (11). In contrast, a subset of CD4^+^ Th1 cells that are highly differentiated (Tbet^+^) and co-expressing KLRG^+^ are present in the lung vasculature and are not efficient in controlling *Mtb* growth (11). Adoptive transfer of the less differentiated CXCR3^+^KLRG^-^CD4^+^ purified parenchymal T cells provided protection upon *Mtb* challenge while the CXCR3^+^KLRG1^+^ T cells are not protective upon transfer. In human studies, CXCR5^+^ CCR5^+^ T cells in the lungs and pleural fluid produced IFN-γ (40–42). In preclinical macaque model of latent and active TB, CXCR3^+^CCR6^+^ co-expressing T cells produced both IL-17 and IFN-γ cytokines in the BAL and were associated with the protective responses in latent TB (43). Finally, in vaccine models of subunit vaccination, CXCR3^+^KLRG^-^ T cells readily trafficked to the lung parenchyma and provided *Mtb* control (37). With the increased resolution provided by our studies using a combination of IT and IV labeling, allow us to further discriminate whether immune cells are localized within the airways or parenchyma or the vascular compartments. Our studies using this new technique show that BCG vaccination induces a mixed population of CXCR3^+^ T cells that are distributed equally between the vasculature, parenchyma and the airway compartments. In sharp contrast, BCG vaccinated hosts that also receive Z-DC transfer show a remarkable enhancement of CXCR3^+^ T cells that localize within the airway and parenchyma compartments.

Our study show that upon BCG vaccination, the primary lung localizing T cells are IL-17 expressing cells that accumulate by day 15, with IFN-γ producing cells accumulating by day 21 post *Mtb* infection. In contrast, BCG vaccinated mice that receive Z-DC transfer recruit Th17 cells by 5 days mostly in the airways and parenchyma followed by IFN-γ-single or IFN-γ/IL-17- coproducing cells accumulating largely in the airways and parenchyma by day 8. Both IFN-γ and IL-17 have varied roles in *Mtb* control during vaccination. In the case of IFN-γ, while recent data has shown that IFN-γ (44) and IFN-γ produced by CD4^+^ T cells (21) are considered redundant, IL-17 is necessary for vaccine-induced control in many models of vaccination (12, 34, 45). More recent work from our lab has also demonstrated a critical role for IL-22 in mediating *Mtb* control. Intriguingly in our model described here, our results show a combined role for IL-17/IL-22 and IFN-γ in conferring early vaccine-induced control of *Mtb* infection. Therefore, it is likely that the mucosal delivery of activated DCs accelerated parenchymal homing of antigen specific CD4^+^ T cell subsets to gain access to the *Mtb*-infected cells in the granuloma and reduce *Mtb* replication through activation of signaling involving both Th1/Th17 cytokine axis.

Upregulation of *Itgae* (Cd103) and Ccr8 genes in CD4^+^ T cells isolated from BCG vaccinated *Mtb*-infected mice that received Z-DC transfer suggest that these molecules may regulate CD4^+^T cell migration and localization within the lung compartments. CD103 is an integrin highly expressed in tissue resident memory T cells (21) and associated with epithelial retention of T cells through binding to E-cadherin expressed by epithelial cells (46). CD103 expressing T cells are present in lung and BAL but absent in blood of *Mtb* infected humans suggesting the fact that CD103 expressing cells are present at the site of *Mtb* infection. CD103 expressing T cells are enriched at the lung parenchyma and airways following mucosal vaccination with BCG or *Bacillus subtilis* spore fusion protein 1 (Spore-FP1) and confers better protection against *Mtb* infection compared to BCG parenteral vaccination (47, 48). Thus, retention of CD103 expressing T cells at the lung interface is likely necessary for providing protection against infection. Deficiency of Cd103 (in mice) correlated with reduced number of mucosal intraepithelial T cells (49). Our results show that despite the increased expression of CD103 on CD4^+^ T cells isolated from BCG vaccinated *Mtb*-infected mice that received Z-DC transfer and localization within the airway and parenchyma, CD103 deficient mice upon BCG vaccination and Z-DC transfer still provided similarly superior protection as BCG vaccinated wild type C57BL/6 mice that received Z-DC transfer. These results suggest redundant features of integrin that may mask the effect of single deficiency of this protein, and compensatory mechanisms are being induced to ensure control of *Mtb* replication in CD103 deficient vaccinated mice that receive Z-DC transfer. Our studies also show for the first time that CCR8 expression is high on CD4^+^ T cells isolated from BCG vaccinated mice receiving Z-DC transfer and expression is highest on airway localized Ag85b-specific CD4^+^ T cells. CCR8 is commonly expressed by T regulatory subsets or T helper type 2 cells for efficient migration of T cell population to the site of inflammation (50). CCL1 was shown to be upregulated upon in vitro infection with *Mtb* and in patients with active tuberculosis versus latently infected controls (51, 52). However, the functional role of CCR8 expressing CD4^+^ T cells during *Mtb* infection remains elusive. Our studies show a functional role for CCR8 expression in the superior protection mediated by the Z-DC transfer in BCG vaccinated mice, as absence of CCR8 expression abrogates Z-DC-mediated protection and accumulation of Ag85B-specific T cells. Together these results suggest that CD103 and CCR8 have pivotal role to play in early CD4^+^ T cell recruitment and localization in the airways and parenchyma to achieving effective control of *Mtb*.

In recent studies, we proposed a role for AMs to localize from airways into the lung parenchyma as an effector mechanism of protection upon *Mtb* infection (10). Consistent with this proposed role for AMs in early *Mtb* control, our new results here demonstrate that during BCG vaccination, AMs continue to be retained within the airways, while in BCG vaccinated hosts which also received Z-DC transfer, AMs are activated rapidly (day 8) and migrate into the parenchyma compartment. In sharp contrast, the AMs in BCG vaccinated mice take up to 20 days to undergo activation. Similarly, recruited macrophages in BCG vaccinated mice are mostly located within the vasculature, while Z-DC transfer activates the RMs to migrate into the airway. These responses appear to be mediated by signaling in epithelial cells as mice deficient in NFκβ signaling in CCSP^+^ epithelial cells abrogate AM accumulation, downstream activation and accumulation of IL-17 producing Ag85B-specific CD4^+^ T cells in the lung. Epithelial cells respond to several external stimulus including IL-17 and IL-22 and activate NFκβ dependent signaling pathways to produce chemokines and other chemotactic factors required to favor other immune cell recruitment and transmigration into inflamed tissues (29, 53). Based on the increased susceptibility of mice lacking NFκβ signaling in epithelial cells compartment, we propose that epithelial cells play an important role in mediating transmigration and specific localization of immune cells within lung parenchyma following being activated by IL-17 and/or IL-22. Lung epithelial cells can produce chemokines such as CXCL9, 10, 11 (54) and 13 in presence of various stimuli (19, 55). Our published data show that IL-17 and IL-22 cytokines are inducers of chemokines following *Mtb* infection (8, 19, 25, 34, 45). Additionally, our published data suggest an important role of the CXCL13/CXCR5 in organizing the iBALT structures which help in rapid containment of the disease (19). Moreover, IL-17 cytokine is also involved in the initial formation of the iBALT structures following *Mtb* infection in mice (26). Therefore, our results show that innate activation of DCs results in activation of epithelial signaling in the lung to amplify accumulation of CD4^+^ T cells that localize within the airways and parenchyma to induce *Mtb* killing of infected macrophages.

In conclusion, using a model of early complete *Mtb* control in BCG vaccinated hosts we show that rapid and early clonal expansion of activated cytokine-producing CD4^+^ T cells in the lung airway and parenchyma compartment are critical for mediating complete and early vaccine-induced protection in *Mtb*-infected mice. Importantly, these protections are driven by early signaling events in the lung epithelial cells that provide the signals required for localization of CD4^+^ T cells within the parenchyma for activation of macrophages, formation of iBALT structures and subsequent *Mtb* killing. Our studies support the emerging idea that Th1/Th17-like activated CD4^+^ T cells are associated with models of sterilizing protection in macaques (23), and in vaccine-induced protection in human TB vaccines M72/AS01E trial (56). Understanding the early immune parameters that mediate effective and early *Mtb* control as demonstrated in this study will shed novel insights into the mechanisms by which vaccine-induced CD4^+^ T cells can be enhanced to mediate complete control of *Mtb*.

## Materials and Methods

### Mice

C57BL/6 (B6), B6.129P2-Il10tm1Cgn/J (Il10^-/-^), B6.129S2(C)-Itgaetm1Cmp/J (*Itgae*^-/-^ or *Cd103*^-/-^) mice were obtained from Jackson Laboratory (Bar Harbor, ME) and bred at Washington University in St. Louis. Cryopreserved sperm from *Ccr8^-^*^/-^ mice were generously donated by Dr. Gwendalyn Randolph from Washington University in St. Louis and the in vitro fertilization was done in the Micro-injection Core at Washington University in St Louis*. Ikk^fl/fl^ Sftpcre* mice were a kind gift from Dr. Pasparakis (University of Cologne). Il-22^-/-*(28)*^ and *Il-17*^-/-(57)^ single knock outs were crossed and bred at Washington University in St. Louis to generate *Il-17/Il22*^-/-^. Mice were age and sex-matched and used between 6-8 weeks of age. All mice were used and housed in accordance with the National Institute of Health guidelines for housing and care of laboratory animals. All the experiments in this study were granted by the Washington University in St Louis Institutional Animal Care and Use Committee under protocol 20160129.

### IFN-**γ** in vivo neutralization

300μg/ml of anti-IFN-γ blocking antibody (Clone XMG1.2, BioXcell) was administered intraperitoneally every other day starting at 8 dpi until the harvest at 20 dpi.

### Bacterial infection and vaccination

*M. bovis* Bacille Calmette–Guerin (BCG Pasteur, Source: Trudeau Institute) and *Mtb* W. Beijing strain, HN878 (BEI Resources) were grown to mid-log phase in Proskauer Beck medium containing 0.05% Tween 80 and frozen in at -80° C. Mice were vaccinated with 1 × 10^6^ colony forming units (CFU) BCG subcutaneous and 4 weeks later infected with ∼100 CFU *Mtb* HN878 via aerosol route using a Glas-Col airborne infection system. At given time points following infection, lungs were collected, homogenized and the tissue homogenates were plated following serial dilutions on 7H11 agar (BD bioscience) to assess bacterial burden (8).

### In vitro culture of BMDCs and transfer

Bone marrow-derived dendritic cells (BMDC) and bone marrow-derived macrophages (BMDMs) were generated as previously described (8). Briefly, cells isolated from the femur and tibia were cultured at 1 × 10^6^ cells/ml in 10 ml of complete DMEM (cDMEM) supplemented with 4% recombinant mouse GM-CSF (Peprotech, Rocky Hill, NJ, USA) at 37°C in 7.5% CO_2_. After 3 days, 10 ml of cDMEM supplemented with 4% mouse GM-CSF was added and incubation continued till day 7. At day 7, non-adherent cells (BMDCs) were collected, counted, plated at 2 × 10^6^ cells/ml in cDMEM and rested overnight at 37°C in 7.5% CO_2_ following which BMDCs were stimulated overnight with Ag85B (20 µg/ml) (New England Peptide) and Zymosan (25 µg/ml) (Sigma) to induce maturation and activation. Mature pulsed BMDCs were collected, washed and 1 × 10^6^ cells in 50 ul PBS were instilled via intratracheal (IT) route at -1 and +4 dpi. For all the adoptive transfer of Z-DCs, Il10^-/-^ BMDC were used.

### Generation of single-cell suspensions from tissues

Lung single-cell suspensions from vaccinated or *Mtb*-infected mice were isolated as previously described (58). Briefly, mice were euthanized with CO_2_ and lungs were perfused with heparin in saline. Lungs were minced and incubated in Collagenase/DNAse for 30 minutes at 37°C. Lung tissue was pushed through a 70 µm nylon screen to obtain a single cell suspension. Red blood cells were lysed and the cells were resuspended in suitable media or buffer for further use.

### CD4^+^ T cell isolation for RNA sequencing and adoptive transfer

Single cells suspensions from infected mice were obtained as before (8). CD4^+^ T cells from differently treated mice were isolated using CD4^+^ microbeads according to manufacturer’s instruction (Miltenyi Biotec). The purity of CD4^+^ T cells was analyzed by flow cytometry after the staining with anti-CD4 antibody and reported to be > 95%. For the RNA sequencing analysis, cells were collected in RLT buffer with β-mercaptoethanol and processed according the manufacturer’s instructions (Qiagen). For T cell transfer, 2 × 10^6^ CD4^+^ T cells were transferred via IT route in PBS into each mouse as previously described.

### RNA-Seq data analysis

Purified mouse lung CD4^+^ T cells were snap-frozen in RLT buffer, and DNase-treated total RNA was extracted using the Qiagen RNeasy Mini kit (Qiagen). RNA-seq libraries were generated using the Clontech SMART-Seq v4 Ultra Low Input RNA Kit for sequencing and the Illumina Nextera XT DNA Library preparation kit following the manufacturer’s protocol. Raw sequencing reads were quality checked for potential sequencing issues and contaminants using FastQC. Adapter sequences, primers, Ns, and reads with quality score below 28 were trimmed using fastq-mcf of ea-utils and PRINSEQ. Reads with a remaining length of less than 20bp after trimming were discarded. Paired end reads were mapped to the mouse genome (mm10) using STAR in a strand specific manner. Read coverage on forward and reverse strands for genome browser visualization was computed using SAMtools, BEDtools, and UCSC Genome Browser utilities. Pairwise differential expression was quantified using DESeq2 (version 1.24.0), with default settings and a 10-5 adjusted P value cutoff for significance, and DESeq2-normalized read counts were used to calculate relative expression (FPKM) values. Heatmap figures were generated in Microsoft Excel, using Z-scores calculated from the FPKM values across all of the samples, individually for each gene. ”Principal Components Analysis (PCA) was performed according to default DESeq2 settings, utilizing the top 500 most variable genes across all samples. Lists of significantly differentially expressed genes were used to test for significant enrichment among KEGG pathways(59) using WebGestalt(60) (default settings, adjusted P = 0.05 threshold for enrichment).

### scRNA-Seq library generation and sequencing

Isolated total lung single cell suspensions were enriched for live cells using dead cell depletion kit according to manufacturer’s instruction (Milteny Biotec) and subjected to droplet-based massively parallel single-cell RNA sequencing using Chromium Single Cell 5’ (v3) Reagent Kit as per manufacturer’s instructions (10x Genomics). Briefly, cell suspensions were loaded at 1,000 cells/µL with the aim to capture 10,000 cells/lane. The 10x Chromium Controller generated GEM droplets, where each cell was labeled with a specific barcode, and each transcript labeled with a unique molecular identifier (UMI) during reverse transcription. The barcoded cDNA was isolated and removed from the BSL-3 space for library generation. The cDNA underwent 11 cycles of amplification, followed by fragmentation, end repair, A-tailing, adapter ligation, and sample index PCR as per the manufacturer’s instructions. Libraries were sequenced on a NovaSeq S4 (200 cycle) flow cell, targeting 50,000 read pairs/cell.

### scRNA-Seq analysis

Sample demultiplexing, barcode processing, and single-cell 5’ counting was performed using the Cell Ranger Single-Cell Software Suite (10x Genomics, version 3). Cell ranger cell count was used to align samples to the reference mm10 genome, quantify and filter reads with a quality score below 30. For TCR, the Seurat package (61) in R was used for subsequent analysis. Cells with mitochondrial content greater than 10% were removed. Filtered data were normalized using a scaling factor of 10,000 nUMI was regressed with a negative binomial model, and data was log transformed. The highly variable genes were selected using the FindVariableFeatures. The principal component analysis was performed using the top 3000 variable genes. Clustering was performed using the FindClusters function. UMAP was used to project cells into two dimensions using 15 first principal components. For T cell re-clustering we chose clusters that were identified as T cells (Cd3d^+^). For these 24758 cells we performed normalization, found variable genes and performed PCA, UMAP and clustering as described above. All visualization was done with ggplot2 R package (62), heatmaps were done with Phantasus website (https://artyomovlab.wustl.edu/phantasus/).

#### Public bulk RNA-seq reanalysis

We re-analyzed the publicly available GSE94964 dataset. With Phantasus, we filtered low expressed genes and did log2(exp+1) and quantile normalization. Further, to get the T resident memory signature for CD4^+^ T cells, we compared CD4^+^ CD69^+^ samples from the lung with CD4^+^CD69^-^ samples from the lung and blood with limma. We have taken 500 upregulated genes. To compare with effector CD4_1 cluster from our data, we run FindMarkers function for CD4_1 cluster versus CD4_2 and CD4 Naïve cells with “MAST” algorithm, logFC threshold of 0.15 and “min.pct” parameter of 0.2. We used the resulted ranked gene list and 500 upregulated genes from the GSE94964 dataset as a signature to run GSEA with the fgsea package.

### Single cell paired TCR**α**/TCR**β** repertoire analysis

Sample demultiplexing and barcode processing was performed using the Cell Ranger Single-Cell Software Suite (10x Genomics). Cell Ranger VDJ v3 was used to align reads to the reference genome (vdj_GRCm38_alts_ensembl) and assemble TCRs. For downstream analysis, only TCRs with 1 productive rearrangement for TCRα chain and 1 productive rearrangement for TCRβ chain were selected. Frequencies of clonotypes were calculated based on number of cells that pass quality control as described above and share both TCRα and TCRβ nucleotide sequences. Gini coefficient was computed with “gini” function from TCR R package. To infer motifs, TCRdist tool was used (63) on all TCRs that satisfied two criteria: that they belonged to CD4 T cells and that they had exactly 1 TCRβ and 1 TCRα chain with CDR3 sequence that passed cellranger quality control. TCRdist was run with default settings for “mouse” organism. We chose three motifs that were most abundant among unique clonotypes, without accounting for clonotype expansion. To characterize the motif and match it to TCRs, we have chosen the most conservative stretches amino acids to represent each motif: “NTGKL” in TCRα and “SLE” in TCRβ for motif 1, “NNNNAP” in TCRα and “E[TR]L” for motif 2, and “NAYKV” in TCRα and “SLE” in TCRβ for motif 3. All clonotypes that contained these sequences were classified as motif-bearing clonotypes. To calculate frequencies of cells for motifs, we calculated how many cells are having motif-bearing TCR clonotype. Further, motif-bearing TCRs were used to visualize motifs by first performing multiple sequence alignment with the msa R package (ClustalW algorithm), and after representing the motifs with the ggseqlogo R package.

### Intratracheal and intravascular staining

Mice were anesthetized with Isoflurane before the IT and IV staining. For IT staining, 0.7 µg/mouse of anti-CD45.2-v500 ab (clone 104, BD Biosciences) in 50 ul of PBS was instilled through intratracheal route 15 minutes before the harvest. For the IV staining, 2.5 µg/mouse of anti-CD45.2-BV605 ab (clone 104, BD) in 100 ul of PBS were injected into the retro-orbital sinus 3 minutes before the harvest using a 26-gauge needle and a tuberculin syringe (10).

### Flow cytometry staining

The following antibodies were from TonBo Biosciences: MHC-II (clone M5/114.15.2), IFN-γ (clone XMG1.2) and CD4 (clone RM4-5). Antibodies purchased from eBioscience (San Diego, CA, USA) were: CD103 (clone 2E7), CD44 (clone IM7) and CXCR3 (clone CXCR3-173). CD11b (clone M1/70), CD11c (clone HL3), Gr1 (clone RB6-8C5), SIGLEC-F (clone E50-2440), CD3 (clone 500A2), CD4 (clone RM4-5) and IL-17 (clone TC11-18H10) were purchased from BD Biosciences. Ag85B tetramers were obtained from NIH tetramer core. For flow cytometric analysis, lung single cell suspensions were stained with tetramer prior to the surface and intracellular staining for 1 hour at 37°C.Intracellular cytokine staining was performed using the BD Cytofix/Cytoperm kit (BD Biosciences) following manufacturer’s instructions. Intracellular staining with anti-IFN-γ and IL-17 was performed for 30 minutes. Cells single stained with each fluorochrome were used as controls for the compensation matrix in the flow cytometry. Samples were acquired on a 4 laser BD Fortessa Flow Cytometer and the analysis was performed using FlowJo (Treestar).

### Immunofluorescence staining

For immunofluorescent staining, formalin fixed and paraffin embedded (FFPE) lung sections were cut, immersed in xylene, and then hydrated in 96% alcohol and phosphate-buffered saline. Antigens were unmasked using a DakoCytomation Target Retrieval Solution (Dako), and non-specific binding was blocked by adding 5% (v/v) normal donkey serum and Fc block (BD). Avidin was used to neutralize endogenous biotin, followed by incubation with biotin (Sigma Aldrich). Sections were then probed with anti-B220 (clone RA3-6B2, BD) and anti-CD3 (clone M-20, Santa Cruz Biotechnology) to detect B cells and T cells respectively. For analysis of B-cell follicles, follicles were outlined with an automated tool of the Zeiss Axioplan 2 microscope (Zeiss), and total area and average size was calculated in squared microns.

### Cytokine and chemokine quantification using Luminex or ELISA

Cytokine and chemokine protein contents in lung homogenates were quantified using Luminex multianalyte technology (Millipore) according to manufacturer’s protocols. IL-17, IFN-γ was quantified by ELISA according to manufacturer’s instructions (R&D).

### Statistical Analysis

The differences between two groups were analyzed using two-tailed student’s t test in Prism 5 (GraphPad). Differences between the means of three or more groups were analyzed using One-way ANOVA with Tukey’s post-test. For comparisons between two or more groups with two independent variables, 2-way ANOVA with Sidak’s or Tukey’s post-test was used. A p-value of <0.05 was considered significant. Raw read counts were used as input for DESeq2(64) (version 1.24.0) differential expression analysis, using default settings and an FDR-adjusted P value threshold of 10^-5^ for significant differential expression. Lists of significantly differentially expressed genes were used to test for significant enrichment among KEGG pathways (59) using WebGestalt (60) (default settings, adjusted P = 0.05 threshold for enrichment).

### Data availability statement

scRNA-Seq data that support the findings of this study have been deposited in GEO (ID GSE150657) and synapse (ID syn22036882). RNA-Seq data that support the findings of this study have been deposited in GEO (accession number: GSE165614). Other data that support the findings of this study are available from the corresponding author upon request.

### Declaration of interest

The authors declare no competing interests.

## Acknowledgement

This work was supported by Washington University School of Medicine, NIH grant HL105427, AI111914-02, AI134236-02 and AI123780 to S.A.K. and D.K., the Department of Molecular Microbiology, Washington University School of Medicine, and Stephen I. Morse Fellowship to S.D. J.R.-M. was supported by funds of the Department of Medicine, University of Rochester, and NIH grant U19 AI91036. We thank the Genome Engineering and iPSC Center and Department of Pathology Micro-Injection Core (Washington University School of Medicine) for assistance in generating the *Ccr8*^-/-^ mice, NIH tetramer core facility for generously providing Ag85B tetramers and Ms. Lan Lu and Ms. Misty Veschak (Washington University School of Medicine) for technical support.

## Author contributions

S.A.K. designed the study, provided funding. S.D., N.D.M., and M.A. performed mouse experiments and compiled the results. E.E., A.S., M.N.A, B.A.R., and M.M. performed the scRNA-Seq and RNA-Seq analysis, comparative transcriptomics, and functional enrichment analysis. J.R.-M. performed histochemical analysis. M.G.N., L.B.B., M.Z., M.N.A., D.K., and S.A.K. interpreted experiments, carried out data analysis and/or provided reagents. S.A.K., S.D., and E.E wrote the manuscript, all authors edited and approved the final version of the manuscript.

## Supplementary Figure legends

**Supplementary Figure 1. Mucosal delivery of Z-DCs in *Mtb*-infected BCG vaccinated mice results in unique *Mtb* antigen specific CD4^+^ T cell activation at the site of infection.** UMAP with CD3^+^ cells, split by condition is shown (**A**). T cell cluster abundances as percent of total CD3^+^ cells across four conditions are shown (**B**). Error bars are mean ± SD for two replicates from each condition. Vac+Z-DC 8 dpi had only one sample. Gene set enrichment analysis (GSEA) shows an enrichment of genes, upregulated in lung Trm (GSE94964), in our comparison between CD4_1 and CD4_2/Naive CD4 cluster (C). Amino acid composition of CDR3α and CDR3β for motif 2 and 3 depicted as sequence logos (**D**). Proportion of CD4^+^ cells, matching motif 1 across all samples and clusters are shown (**E**).

**Supplementary figure 2. RNA-Seq reveals unique gene signatures in CD4^+^ T cells isolated from *Mtb-*infected BCG vaccinated C57BL/6 mice that received Z-DC transfer.** CD4^+^ T cells were isolated from *Mtb-*infected unvaccinated (20 dpi), *Mtb-*infected BCG vaccinated (15 dpi) and *Mtb-*infected BCG vaccinated mice that received Z-DC transfer (8 dpi) and RNA was extracted. Principal component analysis showing genes differentially expressed by CD4^+^ T cells isolated from differently treated mice **(A)**. Table showing top 25 genes differentially expressed in CD4^+^ T by *Mtb-*infected BCG vaccinated mice that received Z-DC transfer cells as compared with *Mtb-*infected BCG vaccinated mice **(B)**. KEGG pathway analysis showing the gene signatures upregulated in CD4^+^ T from by *Mtb-*infected BCG vaccinated mice that received Z-DC transfer cells as compared with *Mtb-*infected BCG vaccinated mice **(C)**. C57BL/6 mice were vaccinated, infected and received Z-DC as described in method. At the time of harvest, the mice were given both anti-CD45.2-v500 and anti-CD45.2-BV605 antibodies through IT and IV route respectively. Lungs were harvested and subjected to flow cytometry. Gating strategy is shown to detect lung T cells populations. T cells were characterized as CD3^+^CD4^+^CD44^hi^Tetramer^+^ (D). These Tetramer^+^ cells were further gated based on CD45.2-v500 and CD45.2-BV605 staining. CD3^+^CD4^+^CD44^hi^Tetramer^+^CD45.2-v500^+^CD45.2-BV605^-^ cells represented the airways populations, CD3^+^CD4^+^CD44^hi^Tetramer^+^CD45.2-v500^-^CD45.2-BV605^+^ represented the vasculature populations and CD3^+^CD4^+^CD44^hi^Tetramer^+^CD45.2-v500^-^CD45.2-BV605^-^ represented the parenchyma populations. These three populations were further analysed for expression of IL-17, IFN-γ, CXCR3, CD103 and CCR8.

**Supplementary figure 3. Mucosal delivery of Z-DCs induces CD4^+^ cell activation through epithelial signaling for vaccine-induced immunity.** C57BL/6 mice were vaccinated, infected and received Z-DC as described in method. At the time of harvest, the mice were given both anti-CD45.2-v500 and anti-CD45.2-BV605 antibodies through IT and IV route respectively. Lungs were harvested and subjected to flow cytometry. The number of CD4^+^CD44^hi^ T cells were detected by flow cytometry (**A**). The distribution of CD4^+^CD44^hi^ T cells (**B**) in *Mtb*-infected BCG vaccinated mice and *Mtb*-infected BCG vaccinated mice that received Z-DC transfer were measured in lung airways (red bar), parenchyma (blue bar) and vasculature (green bar) regions by flow cytometry. n =4-5 mice per group. Gating strategy is shown to detect myeloid cell populations (**C**). AMs were characterised as CD11C^+^CD11B^-^SiglecF^+^, RMs were characterised as CD11C^-^CD11B^+^Gr1^-^. AMs and RMs were further characterised based on the expression of CD45.2-v500 and CD45.2-BV605 to determine their location as mentioned above. B6, and *Cd103^-/-^* mice were vaccinated, infected and received Z-DC as described in method. Mice were harvested at 20 dpi and lung bacterial burden was determined by plating **(D)**. n =4-5 mice per group. In a separate experiment, *Ikk^fl/fl^Sftpcre* mice were vaccinated, infected and received Z-DC as described in method. To track the immune cells mice were given anti-CD45.2-v500 and anti- CD45.2-BV605 antibodies as before. Lungs were harvested at 20 dpi and MHC-II MFI (mean fluorescent intensity-MFI) on RMs (**E**), the total numbers of CD4^+^CD44^hi^ (**F**), CD4^+^CD44^hi^TET^+^ T (**G**) cells were determined by flow cytometry (red=*Ikk^fl/fl^*, blue= *Ikk^fl/fl^Sftpcre*). n=4-5 mice per group. Data represented as mean ± SD. ** p≤ 0.01, *** p≤0.001, ****p≤ 0.0001 either by two way ANOVA (**A and B**), or Student’s t test (actual p values are shown) (**D-G**).

## Supplementary Table legends

Table S1: Description of Cell proportion in each cluster per condition from scRNA-seq analysis.

Table S2: Description of TCRs per condition from scRNA-seq analysis.

## Notes

### Competing Interest Statement

The authors have declared no competing interest.

## References

1. WHO. 2018. Tuberculosis Fact sheet http://www.who.int/mediacentre/factsheets/fs104/en/. Accessed

2. Tait DR, Hatherill M, Van Der Meeren O, Ginsberg AM, Van Brakel E, Salaun B, Scriba TJ, Akite EJ, Ayles HM, Bollaerts A, Demoitie MA, Diacon A, Evans TG, Gillard P, Hellstrom E, Innes JC, Lempicki M, Malahleha M, Martinson N, Mesia Vela D, Muyoyeta M, Nduba V, Pascal TG, Tameris M, Thienemann F, Wilkinson RJ, Roman F. 2019. Final Analysis of a Trial of M72/AS01E Vaccine to Prevent Tuberculosis. N Engl J Med 381:2429–2439.

3. Darrah PA, Zeppa JJ, Maiello P, Hackney JA, Wadsworth MH, 2nd, Hughes TK, Pokkali S, Swanson PA, 2nd, Grant NL, Rodgers MA, Kamath M, Causgrove CM, Laddy DJ, Bonavia A, Casimiro D, Lin PL, Klein E, White AG, Scanga CA, Shalek AK, Roederer M, Flynn JL, Seder RA. 2020. Prevention of tuberculosis in macaques after intravenous BCG immunization. Nature 577:95–102.

4. Kaushal D, Foreman TW, Gautam US, Alvarez X, Adekambi T, Rangel-Moreno J, Golden NA, Johnson AM, Phillips BL, Ahsan MH, Russell-Lodrigue KE, Doyle LA, Roy CJ, Didier PJ, Blanchard JL, Rengarajan J, Lackner AA, Khader SA, Mehra S. 2015. Mucosal vaccination with attenuated Mycobacterium tuberculosis induces strong central memory responses and protects against tuberculosis. Nat Commun 6:8533.

5. Verreck FAW, Tchilian EZ, Vervenne RAW, Sombroek CC, Kondova I, Eissen OA, Sommandas V, van der Werff NM, Verschoor E, Braskamp G, Bakker J, Langermans JAM, Heidt PJ, Ottenhoff THM, van Kralingen KW, Thomas AW, Beverley PCL, Kocken CHM. 2017. Variable BCG efficacy in rhesus populations: Pulmonary BCG provides protection where standard intra-dermal vaccination fails. Tuberculosis (Edinb) 104:46–57.

6. Caruso AM, Serbina N, Klein E, Triebold K, Bloom BR, Flynn JL. 1999. Mice deficient in CD4 T cells have only transiently diminished levels of IFN-gamma, yet succumb to tuberculosis. J Immunol 162:5407–5416.

7. Urdahl KB, Shafiani S, Ernst JD. 2011. Initiation and regulation of T-cell responses in tuberculosis. Mucosal immunology 4:288–293.

8. Griffiths KL, Ahmed M, Das S, Gopal R, Horne W, Connell TD, Moynihan KD, Kolls JK, Irvine DJ, Artyomov MN, Rangel-Moreno J, Khader SA. 2016. Targeting dendritic cells to accelerate T-cell activation overcomes a bottleneck in tuberculosis vaccine efficacy. Nat Commun 7:13894.

9. Portal-Celhay C, Tufariello JM, Srivastava S, Zahra A, Klevorn T, Grace PS, Mehra A, Park HS, Ernst JD, Jacobs WR, Jr., Philips JA. 2016. Mycobacterium tuberculosis EsxH inhibits ESCRT-dependent CD4(+) T-cell activation. Nat Microbiol 2:16232.

10. Dunlap MD, Howard N, Das S, Scott N, Ahmed M, Prince O, Rangel-Moreno J, Rosa BA, Martin J, Kaushal D, Kaplan G, Mitreva M, Kim KW, Randolph GJ, Khader SA. 2018. A novel role for C-C motif chemokine receptor 2 during infection with hypervirulent Mycobacterium tuberculosis. Mucosal Immunol 11:1727–1742.

11. Sakai S, Kauffman KD, Schenkel JM, McBerry CC, Mayer-Barber KD, Masopust D, Barber DL. 2014. Cutting edge: control of Mycobacterium tuberculosis infection by a subset of lung parenchyma-homing CD4 T cells. J Immunol 192:2965–2969.

12. Khader SA, Bell GK, Pearl JE, Fountain JJ, Rangel-Moreno J, Cilley GE, Shen F, Eaton SM, Gaffen SL, Swain SL, Locksley RM, Haynes L, Randall TD, Cooper AM. 2007. IL-23 and IL-17 in the establishment of protective pulmonary CD4+ T cell responses after vaccination and during Mycobacterium tuberculosis challenge. Nat Immunol 8:369–377.

13. Ogongo P, Porterfield JZ, Leslie A. 2019. Lung Tissue Resident Memory T-Cells in the Immune Response to Mycobacterium tuberculosis. Front Immunol 10:992.

14. Kumar BV, Ma W, Miron M, Granot T, Guyer RS, Carpenter DJ, Senda T, Sun X, Ho SH, Lerner H, Friedman AL, Shen Y, Farber DL. 2017. Human Tissue-Resident Memory T Cells Are Defined by Core Transcriptional and Functional Signatures in Lymphoid and Mucosal Sites. Cell Rep 20:2921–2934.

15. Karlsson AK, Walles K, Bladh H, Connolly S, Skrinjar M, Rosendahl A. 2011. Small molecule antagonists of CCR8 inhibit eosinophil and T cell migration. Biochem Biophys Res Commun 407:764–771.

16. Mami-Chouaib F, Blanc C, Corgnac S, Hans S, Malenica I, Granier C, Tihy I, Tartour E. 2018. Resident memory T cells, critical components in tumor immunology. J Immunother Cancer 6:87.

17. Hara-Chikuma M, Chikuma S, Sugiyama Y, Kabashima K, Verkman AS, Inoue S, Miyachi Y. 2012. Chemokine-dependent T cell migration requires aquaporin-3-mediated hydrogen peroxide uptake. J Exp Med 209:1743–1752.

18. Lai EC. 2004. Notch signaling: control of cell communication and cell fate. Development 131:965–973.

19. Slight SR, Rangel-Moreno J, Gopal R, Lin Y, Fallert Junecko BA, Mehra S, Selman M, Becerril-Villanueva E, Baquera-Heredia J, Pavon L, Kaushal D, Reinhart TA, Randall TD, Khader SA. 2013. CXCR5+ T helper cells mediate protective immunity against tuberculosis. J Clin Invest 123:712–726.

20. Lindenstrom T, Moguche A, Damborg M, Agger EM, Urdahl K, Andersen P. 2018. T Cells Primed by Live Mycobacteria Versus a Tuberculosis Subunit Vaccine Exhibit Distinct Functional Properties. EBioMedicine 27:27–39.

21. Sallin MA, Sakai S, Kauffman KD, Young HA, Zhu J, Barber DL. 2017. Th1 Differentiation Drives the Accumulation of Intravascular, Non-protective CD4 T Cells during Tuberculosis. Cell Rep 18:3091–3104.

22. Gopal R, Monin L, Slight S, Uche U, Blanchard E, Fallert Junecko BA, Ramos-Payan R, Stallings CL, Reinhart TA, Kolls JK, Kaushal D, Nagarajan U, Rangel-Moreno J, Khader SA. 2014. Unexpected role for IL-17 in protective immunity against hypervirulent Mycobacterium tuberculosis HN878 infection. PLoS Pathog 10:e1004099.

23. Dijkman K, Sombroek CC, Vervenne RAW, Hofman SO, Boot C, Remarque EJ, Kocken CHM, Ottenhoff THM, Kondova I, Khayum MA, Haanstra KG. 2019. Prevention of tuberculosis infection and disease by local BCG in repeatedly exposed rhesus macaques. 25:255–262.

24. McAllister F, Henry A, Kreindler JL, Dubin PJ, Ulrich L, Steele C, Finder JD, Pilewski JM, Carreno BM, Goldman SJ, Pirhonen J, Kolls JK. 2005. Role of IL-17A, IL-17F, and the IL-17 receptor in regulating growth-related oncogene-alpha and granulocyte colony-stimulating factor in bronchial epithelium: implications for airway inflammation in cystic fibrosis. J Immunol 175:404–412.

25. Khader SA, Guglani L, Rangel-Moreno J, Gopal R, Junecko BA, Fountain JJ, Martino C, Pearl JE, Tighe M, Lin YY, Slight S, Kolls JK, Reinhart TA, Randall TD, Cooper AM. 2011. IL-23 is required for long-term control of Mycobacterium tuberculosis and B cell follicle formation in the infected lung. J Immunol 187:5402–5407.

26. Rangel-Moreno J, Carragher DM, de la Luz Garcia-Hernandez M, Hwang JY, Kusser K, Hartson L, Kolls JK, Khader SA, Randall TD. 2011. The development of inducible bronchus-associated lymphoid tissue depends on IL-17. Nat Immunol 12:639–646.

27. Kulkarni N, Pathak M, Lal G. 2017. Role of chemokine receptors and intestinal epithelial cells in the mucosal inflammation and tolerance. J Leukoc Biol 101:377–394.

28. Treerat P, Prince O, Cruz-Lagunas A, Munoz-Torrico M, Salazar-Lezama MA, Selman M, Fallert-Junecko B, Reinhardt TA, Alcorn JF, Kaushal D, Zuniga J, Rangel-Moreno J, Kolls JK, Khader SA. 2017. Novel role for IL-22 in protection during chronic Mycobacterium tuberculosis HN878 infection. Mucosal Immunol 10:1069–1081.

29. Hernandez-Santos N, Wiesner DL, Fites JS, McDermott AJ, Warner T, Wuthrich M, Klein BS. 2018. Lung Epithelial Cells Coordinate Innate Lymphocytes and Immunity against Pulmonary Fungal Infection. Cell Host Microbe 23:511–522.e515.

30. Perez-Nazario N, Rangel-Moreno J, O’Reilly MA, Pasparakis M, Gigliotti F, Wright TW. 2013. Selective ablation of lung epithelial IKK2 impairs pulmonary Th17 responses and delays the clearance of Pneumocystis. J Immunol 191:4720–4730.

31. Harding CV, Boom WH. 2010. Regulation of antigen presentation by Mycobacterium tuberculosis: a role for Toll-like receptors. Nat Rev Microbiol 8:296–307.

32. Carpenter SM, Yang JD. 2017. Vaccine-elicited memory CD4+ T cell expansion is impaired in the lungs during tuberculosis. 13:e1006704.

33. Hope JC, Kwong LS, Sopp P, Collins RA, Howard CJ. 2000. Dendritic cells induce CD4+ and CD8+ T-cell responses to Mycobacterium bovis and M. avium antigens in Bacille Calmette Guérin vaccinated and nonvaccinated cattle. Scand J Immunol 52:285–291.

34. Gopal R, Lin Y, Obermajer N, Slight S, Nuthalapati N, Ahmed M, Kalinski P, Khader SA. 2012. IL-23-dependent IL-17 drives Th1-cell responses following Mycobacterium bovis BCG vaccination. Eur J Immunol 42:364–373.

35. Moguche AO, Musvosvi M, Penn-Nicholson A, Plumlee CR, Mearns H, Geldenhuys H, Smit E, Abrahams D, Rozot V, Dintwe O, Hoff ST, Kromann I, Ruhwald M, Bang P, Larson RP, Shafiani S, Ma S, Sherman DR, Sette A, Lindestam Arlehamn CS, McKinney DM, Maecker H, Hanekom WA, Hatherill M, Andersen P, Scriba TJ, Urdahl KB. 2017. Antigen Availability Shapes T Cell Differentiation and Function during Tuberculosis. Cell Host Microbe 21:695–706.e695.

36. Lin PL, Dietrich J, Tan E, Abalos RM, Burgos J, Bigbee C, Bigbee M, Milk L, Gideon HP, Rodgers M, Cochran C, Guinn KM, Sherman DR, Klein E, Janssen C, Flynn JL, Andersen P. 2012. The multistage vaccine H56 boosts the effects of BCG to protect cynomolgus macaques against active tuberculosis and reactivation of latent Mycobacterium tuberculosis infection. J Clin Invest 122:303–314.

37. Woodworth JS, Cohen SB, Moguche AO, Plumlee CR, Agger EM, Urdahl KB, Andersen P. 2017. Subunit vaccine H56/CAF01 induces a population of circulating CD4 T cells that traffic into the Mycobacterium tuberculosis-infected lung. Mucosal Immunol 10:555–564.

38. Carpenter SM, Yang JD, Lee J, Barreira-Silva P, Behar SM. 2017. Vaccine-elicited memory CD4+ T cell expansion is impaired in the lungs during tuberculosis. PLoS Pathog 13:e1006704.

39. Srivastava S, Ernst JD. 2013. Cutting edge: Direct recognition of infected cells by CD4 T cells is required for control of intracellular Mycobacterium tuberculosis in vivo. J Immunol 191:1016–1020.

40. Saha PK, Sharma PK, Sharma SK, Singh A, Mitra DK. 2013. Recruitment of Th1 effector cells in human tuberculosis: hierarchy of chemokine receptor(s) and their ligands. Cytokine 63:43–51.

41. Perreau M, Rozot V, Welles HC, Belluti-Enders F, Vigano S, Maillard M, Dorta G, Mazza-Stalder J, Bart PA, Roger T, Calandra T, Nicod L, Harari A. 2013. Lack of Mycobacterium tuberculosis-specific interleukin-17A-producing CD4+ T cells in active disease. Eur J Immunol 43:939–948.

42. Lindestam Arlehamn CS, Gerasimova A, Mele F, Henderson R, Swann J, Greenbaum JA, Kim Y, Sidney J, James EA, Taplitz R, McKinney DM, Kwok WW, Grey H, Sallusto F, Peters B, Sette A. 2013. Memory T cells in latent Mycobacterium tuberculosis infection are directed against three antigenic islands and largely contained in a CXCR3+CCR6+ Th1 subset. PLoS Pathog 9:e1003130.

43. Shanmugasundaram U, Bucsan AN, Ganatra SR, Ibegbu C, Quezada M, Blair RV, Alvarez X, Velu V, Kaushal D, Rengarajan J. 2020. Pulmonary Mycobacterium tuberculosis control associates with CXCR3- and CCR6-expressing antigen-specific Th1 and Th17 cell recruitment. JCI Insight 5.

44. Gopal R, Monin L, Torres D, Slight S, Mehra S, McKenna KC, Fallert Junecko BA, Reinhart TA, Kolls J, Baez-Saldana R, Cruz-Lagunas A, Rodriguez-Reyna TS, Kumar NP, Tessier P, Roth J, Selman M, Becerril-Villanueva E, Baquera-Heredia J, Cumming B, Kasprowicz VO, Steyn AJ, Babu S, Kaushal D, Zuniga J, Vogl T, Rangel-Moreno J, Khader SA. 2013. S100A8/A9 proteins mediate neutrophilic inflammation and lung pathology during tuberculosis. Am J Respir Crit Care Med 188:1137–1146.

45. Kumar P, Monin L, Castillo P, Elsegeiny W, Horne W, Eddens T, Vikram A, Good M, Schoenborn AA, Bibby K, Montelaro RC, Metzger DW, Gulati AS, Kolls JK. 2016. Intestinal Interleukin-17 Receptor Signaling Mediates Reciprocal Control of the Gut Microbiota and Autoimmune Inflammation. Immunity 44:659–671.

46. Cepek KL, Shaw SK, Parker CM, Russell GJ, Morrow JS, Rimm DL, Brenner MB. 1994. Adhesion between epithelial cells and T lymphocytes mediated by E-cadherin and the alpha E beta 7 integrin. Nature 372:190–193.

47. Perdomo C, Zedler U, Kuhl AA, Lozza L, Saikali P, Sander LE, Vogelzang A, Kaufmann SH, Kupz A. 2016. Mucosal BCG Vaccination Induces Protective Lung-Resident Memory T Cell Populations against Tuberculosis. MBio 7.

48. Copland A, Diogo GR, Hart P, Harris S, Tran AC, Paul MJ, Singh M, Cutting SM, Reljic R. 2018. Mucosal Delivery of Fusion Proteins with Bacillus subtilis Spores Enhances Protection against Tuberculosis by Bacillus Calmette-Guérin. Front Immunol 9:346.

49. Schön MP, Arya A, Murphy EA, Adams CM, Strauch UG, Agace WW, Marsal J, Donohue JP, Her H, Beier DR, Olson S, Lefrancois L, Brenner MB, Grusby MJ, Parker CM. 1999. Mucosal T lymphocyte numbers are selectively reduced in integrin alpha E (CD103)-deficient mice. J Immunol 162:6641–6649.

50. Soler D, Chapman TR, Poisson LR, Wang L, Cote-Sierra J, Ryan M, McDonald A, Badola S, Fedyk E, Coyle AJ, Hodge MR, Kolbeck R. 2006. CCR8 expression identifies CD4 memory T cells enriched for FOXP3+ regulatory and Th2 effector lymphocytes. J Immunol 177:6940–6951.

51. Thuong NT, Dunstan SJ, Chau TT, Thorsson V, Simmons CP, Quyen NT, Thwaites GE, Thi Ngoc Lan N, Hibberd M, Teo YY, Seielstad M, Aderem A, Farrar JJ, Hawn TR. 2008. Identification of tuberculosis susceptibility genes with human macrophage gene expression profiles. PLoS Pathog 4:e1000229.

52. Blischak JD, Tailleux L, Myrthil M, Charlois C, Bergot E, Dinh A, Morizot G, Chény O, Platen CV, Herrmann JL. 2017. Predicting susceptibility to tuberculosis based on gene expression profiling in dendritic cells. 7:5702.

53. Tully JE, Nolin JD, Guala AS, Hoffman SM, Roberson EC, Lahue KG, van der Velden J, Anathy V, Blackwell TS, Janssen-Heininger YM. 2012. Cooperation between classical and alternative NF-κB pathways regulates proinflammatory responses in epithelial cells. Am J Respir Cell Mol Biol 47:497–508.

54. Sauty A, Dziejman M, Taha RA, Iarossi AS, Neote K, Garcia-Zepeda EA, Hamid Q, Luster AD. 1999. The T cell-specific CXC chemokines IP-10, Mig, and I-TAC are expressed by activated human bronchial epithelial cells. J Immunol 162:3549–3558.

55. Wang GZ, Cheng X, Zhou B, Wen ZS, Huang YC, Chen HB, Li GF, Huang ZL, Zhou YC, Feng L, Wei MM, Qu LW, Cao Y, Zhou GB. 2015. The chemokine CXCL13 in lung cancers associated with environmental polycyclic aromatic hydrocarbons pollution. Elife 4.

56. Sable SB, Posey JE, Scriba TJ. 2019. Tuberculosis Vaccine Development: Progress in Clinical Evaluation. Clin Microbiol Rev 33.

57. Gopal R, Monin L, Slight S, Uche U, Blanchard E, Fallert Junecko BA, Ramos-Payan R, Stallings CL, Reinhart TA, Kolls JK, Kaushal D, Nagarajan U, Rangel-Moreno J, Khader SA. 2014. Unexpected role for IL-17 in protective immunity against hypervirulent Mycobacterium tuberculosis HN878 infection.

58. Ardain A, Domingo-Gonzalez R, Das S, Kazer SW, Howard NC, Singh A, Ahmed M, Nhamoyebonde S, Rangel-Moreno J, Ogongo P, Lu L, Ramsuran D, de la Luz Garcia-Hernandez M, T KU, Darby M, Park E, Karim F, Melocchi L, Madansein R, Dullabh KJ, Dunlap M, Marin-Agudelo N, Ebihara T, Ndung’u T, Kaushal D, Pym AS, Kolls JK, Steyn A, Zuniga J, Horsnell W, Yokoyama WM, Shalek AK, Kloverpris HN, Colonna M, Leslie A, Khader SA. 2019. Group 3 innate lymphoid cells mediate early protective immunity against tuberculosis. Nature 570:528–532.

59. Kanehisa M, Sato Y, Furumichi M, Morishima K, Tanabe M. 2019. New approach for understanding genome variations in KEGG. Nucleic Acids Res 47:D590–d595.

60. Liao Y, Wang J, Jaehnig EJ, Shi Z, Zhang B. 2019. WebGestalt 2019: gene set analysis toolkit with revamped UIs and APIs. Nucleic Acids Res 47:W199–w205.

61. Butler A, Hoffman P, Smibert P, Papalexi E, Satija R. 2018. Integrating single-cell transcriptomic data across different conditions, technologies, and species. Nat Biotechnol 36:411–420.

62. Wickham H. 2016. ggplot2: Elegant Graphics for Data Analysis. Springer-Verlag New York.

63. Dash P, Fiore-Gartland AJ, Hertz T, Wang GC, Sharma S, Souquette A, Crawford JC, Clemens EB, Nguyen THO, Kedzierska K, La Gruta NL, Bradley P, Thomas PG. 2017. Quantifiable predictive features define epitope-specific T cell receptor repertoires. Nature 547:89–93.

64. Love MI, Huber W, Anders S. 2014. Moderated estimation of fold change and dispersion for RNA-seq data with DESeq2. Genome Biol 15:550.

