## Supplementary figures and images for "Lung epithelial signaling mediates early vaccine-induced CD4^+^ T cell activation and *Mtb* control"

### Supplemental Figure 1

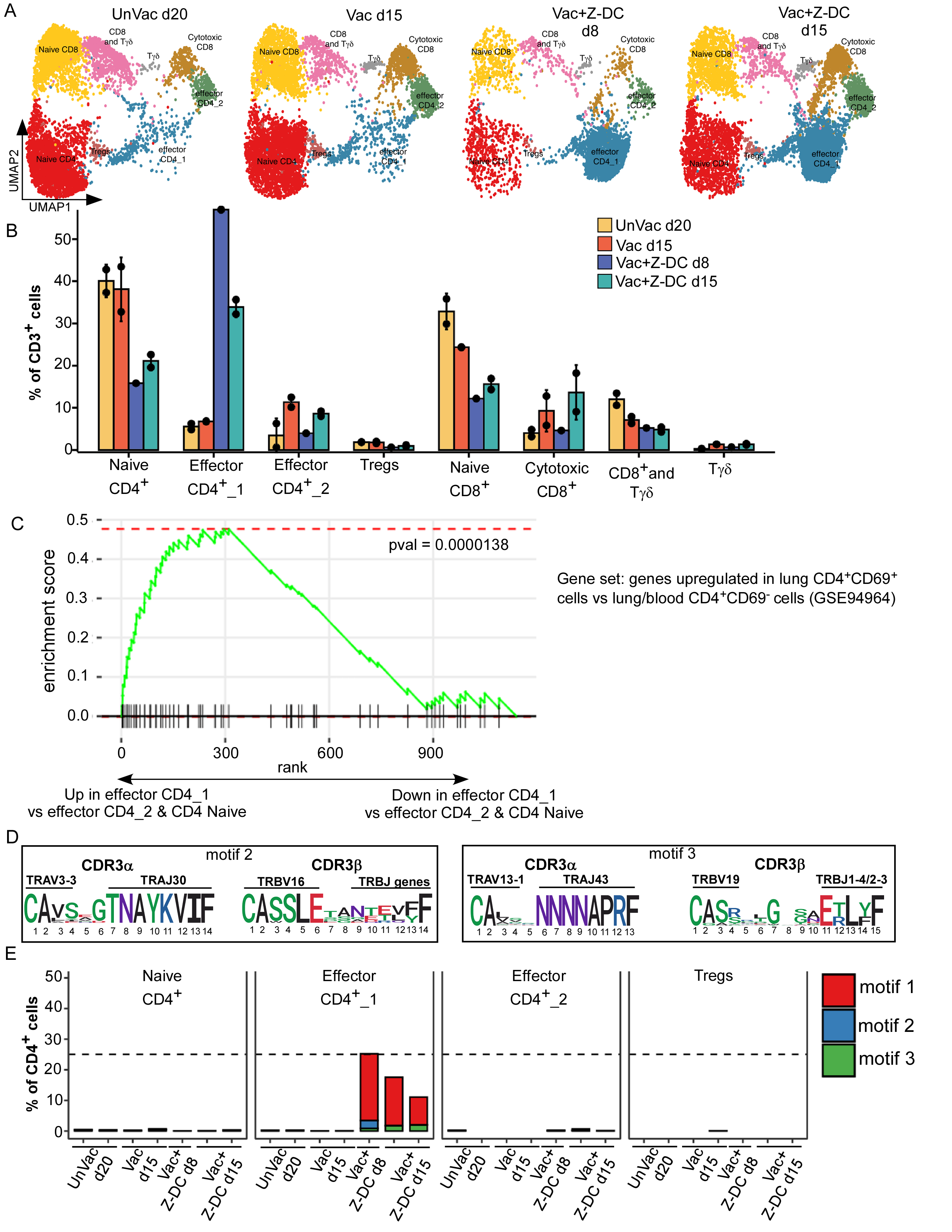

### Supplemental Figure 2

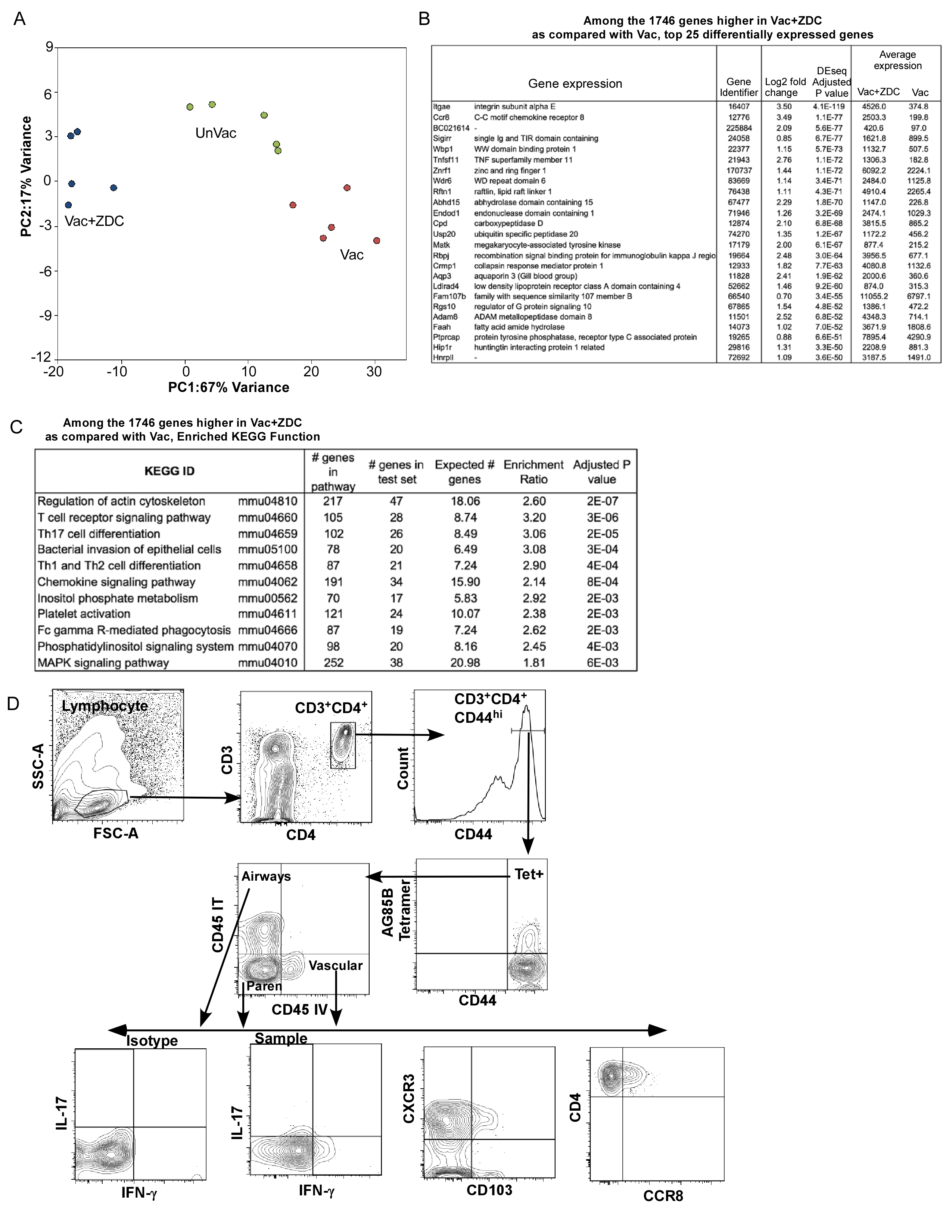

### Supplemental Figure 3

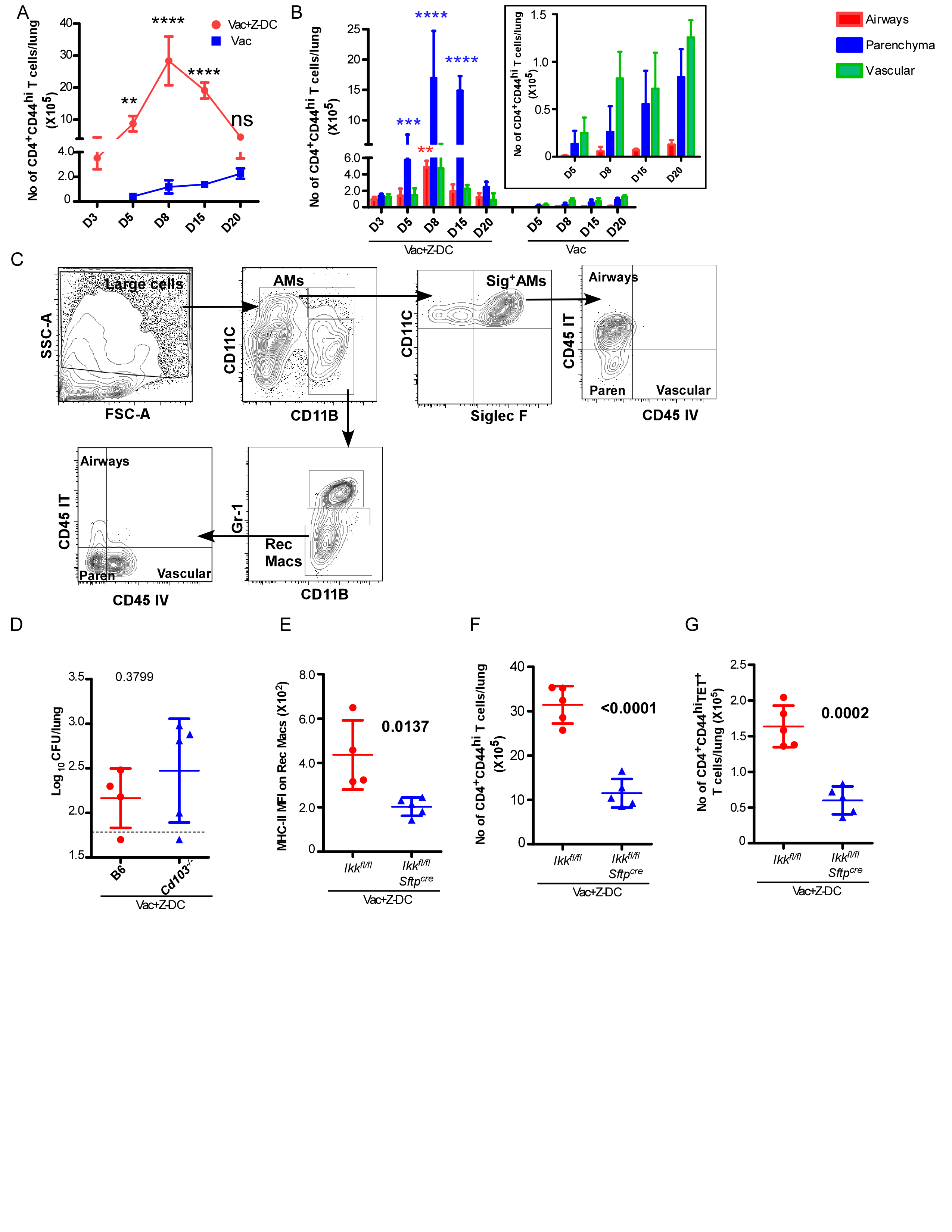
